# Single cell analysis of quiescent HIV infection reveals host transcriptional profiles that regulate proviral latency

**DOI:** 10.1101/303198

**Authors:** Todd Bradley, Guido Ferrari, Barton F Haynes, David M Margolis, Edward P Browne

## Abstract

The latent HIV reservoir is diverse, but most studies of HIV latency have used bulk cell assays. Here we characterized cell line and primary cell models of HIV latency with single cell qPCR (sc-qPCR) for viral RNA (vRNA), and single cell RNAseq (scRNAseq). sc-qPCR revealed distinct populations of cells transcribing vRNA across a wide range of levels. Strikingly, scRNAseq of latently infected primary cells revealed a relationship between vRNA levels and the transcriptomic profiles within the population. Cells with the greatest level of HIV silencing expressed a specific set of host genes including markers of central memory T cells. By contrast, latently infected cells with higher levels of HIV transcription expressed markers of activated and effector T cells. These data reveal that heterogeneous behaviors of HIV proviruses within the latent reservoir are influenced by the host cell transcriptional program. Therapeutic modulation of these programs may reverse or enforce HIV latency.

## Introduction

Persistent HIV infection in the presence of antiretroviral therapy is characterized in part by transcriptional silencing of the virus in latently infected cells. The mechanisms that enforce latency are complex and only partially understood, but involve a repressive chromatin state that is regulated by diverse histone modifications (Turner and Margolis, 2017). Such latently infected cells appear to persist in infected patients for decades, their frequency little changed by years of antiretroviral therapy (ART) (Andrade et al., 2013; Palmer et al., 2008). Furthermore, latently infected CD4+ T cells can contribute to the rebound of infection upon cessation of antiviral therapy, and thus are a principal barrier to curing HIV infection (Chun et al., 2010; Davey et al., 1999; Finzi et al., 1997). As such, intensive efforts currently focus on strategies to eliminate HIV infected reservoir cells. A major strategy to achieve this goal involves pharmacological treatment with latency reversing agents (LRAs) to upregulate HIV expression in infected cells, such that these cells may then be cleared (Archin et al., 2012; Søgaard et al., 2015; Zhu et al., 2012). However, on the level of viral transcription, LRAs are thus far ineffective in the majority of latently infected cells (Ho et al., 2013). The reasons for this inefficiency are unclear. Multiple restrictions of proviral expression may limit the response to single LRAs, as may the heterogeneous nature of latently infected cells themselves. In particular, the role of the host cell’s transcriptional environment in the establishment of latency is an area that has been little explored. Although several host factors, such as CDK9 and CyclinT1, are known to regulate HIV transcription in various models of latency (Friedman et al., 2011; Tripathy et al., 2015; Tyagi et al., 2010), and epigenetic features play a central role in antagonizing or augmenting the role of viral transactivation (He et al., 2002; Turner and Margolis, 2017; Van Lint et al., 2013), it has also been demonstrated that establishment of latency can be driven by stochastic fluctuations in the viral transcription factor Tat (Razooky et al., 2015). As such, the extent to which the host cell’s intracellular milieu impacts this process is still unclear.

The deficiency in our understanding of these questions derives from the majority of studies of HIV-1 latency having only considered cells as populations, studied in bulk culture, and thus crucial information about the behavior of individual cells has not been obtained. The latently infected CD4+ T cell reservoir is inherently diverse, with each provirus exhibiting a potentially unique combination of the effects of integration site, epigenetic modifications, and infected cell phenotype. For example, CD4 T cells, the major host cell for HIV infection, can exist as several different developmental stages, categorized as naïve (Tn) central (Tcm) and effector memory (Tem), and effector cells. Each of these subtypes have distinct transcriptional and epigenetic programs that could impact the activity of the integrated HIV promoter (Durek et al., 2016). Additionally, biological noise and stochastic fluctuations in transcriptional activity could play an important role in either establishment or reversal of latency (Dar et al., 2014).

Given the multitude of factors that can impact the establishment and maintenance of latent infection, the application of methods that permit the analysis of single cells will be required to fully characterize latency and the mechanisms that determine HIV latency. Recent technological breakthroughs in analysis of individual cells by RNAseq (scRNAseq) now permit detailed characterization of heterogeneous behaviors of individual cells (Hashimshony et al., 2016; Macosko et al., 2015; Picelli et al., 2013; Villani and Shekhar, 2017). These methods have provided novel insights into biological systems and revealed surprising diversity in cultures of cells previously assumed to be uniform (Buettner et al., 2015; Shalek et al., 2014; Villani and Shekhar, 2017). We hypothesize that, within the latently infected population, there exist subpopulations with differing patterns of viral and host gene transcription at rest and after host cell activation. Furthermore, we hypothesize that this transcriptional diversity within latently infected cell populations originates from the actions of an identifiable set of host genes. To investigate these hypotheses, we applied two single cell assays to models of latent HIV infection. Analysis of vRNA at the single cell level revealed the existence of diverse levels of vRNA expression both at rest and after LRA stimulation. Furthermore scRNAseq of latently infected primary cells indicated that the level of HIV gene expression was correlated with a specific set of host cell genes. This latency associated signature reveals that silencing of HIV in primary cells is regulated by the underlying transcriptional program of the infected cells. These insights illustrate an important role for the host cell environment in HIV latency, and will guide the development of therapies that can achieve optimal reactivation of the latent reservoir.

## Results

### In a cell line latency model, viral RNA induction by LRAs is heterogeneous, and a threshold effect is seen before viral protein expression

To measure viral RNA (vRNA) in individual latently infected CD4+ T cells, rather than on a population level, we used an approach that combined flow sorting of single cells into 96-well PCR plates followed by quantitative real-time PCR for unspliced HIV RNA. We first analyzed N6 cells, a Jurkat derived CD4+ T cell line that is latently infected with HIV. This cell line contains a full-length integrated copy of the NL4-3 strain of HIV, with the *nef* open reading frame replaced by coding sequence for the murine Heat-shock antigen (HSA) reporter. Flow cytometry for HSA protein expression thus allows us to identify HIV reactivation in N6 cells. We stimulated N6 cells with three different latency reversing agents - the histone deacetylase inhibitor vorinostat (3uM), the protein kinase C (PKC) agonist prostratin (3uM), or tumor necrosis factor-alpha (TNFα; 100ng/mL) and assayed vRNA levels in 144 cells for each condition at 24 hr. (Figure 1A). In unstimulated cells, the majority of cells had undetectable vRNA levels, but a sub-population (19%) expressed low levels of vRNA, detectable but typically below the lower limit of quantification. Upon stimulation, cells treated with either of the three LRAs exhibited a strong upregulation of vRNA and the appearance of detectable HSA protein expression. A wide range of vRNA levels from 1 to 3442 copies per cell was detected across the population, with coefficients of variation ranging from 93% for vorinostat, to 164% for TNFα,. vRNA levels did not exceed 3442 copies per cell, suggesting a uniform restriction of vRNA across the clonal cell line. Notably, expression of virally encoded HSA became pronounced only when vRNA levels were above 500 copies per cell, suggesting that a threshold of vRNA is required for translation of a detectable quantity of HSA in these cells. For all LRAs, the percent of vRNA+ cells exceeded the percent of HSA+ cells, indicating that protein-based viral reporters significantly underestimate the fraction of responding cells (Figure 1B). Thus, these data demonstrate that stimulation of latently infected cells with LRAs induces a broad spectrum of diverse vRNA/antigen responses, and that a significant population of vRNA+ cells can be detected that are not producing detectable viral antigen.

**Fig 1.**
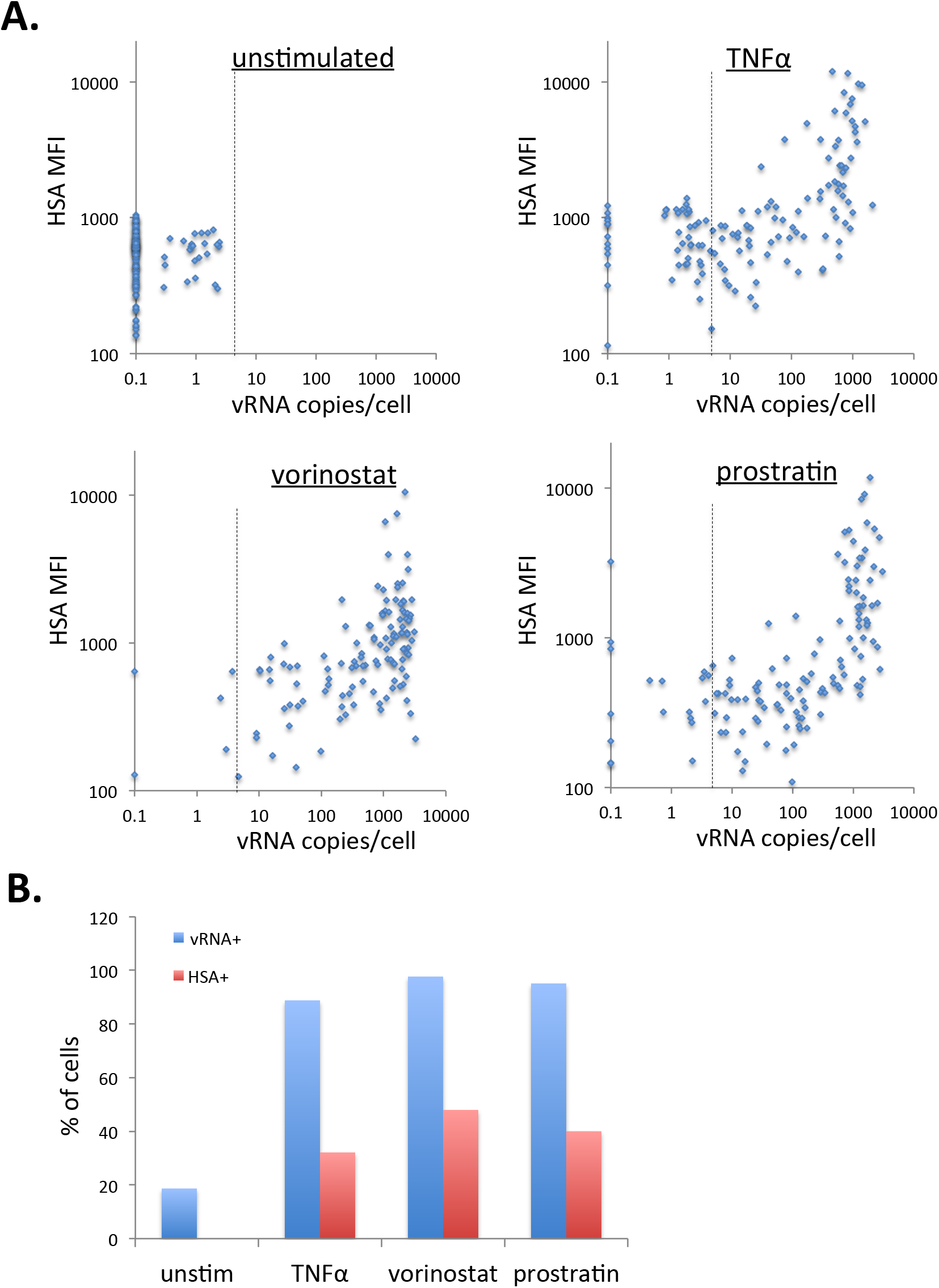
In a cell line latency model, viral RNA induction by LRAs is heterogeneous, and a threshold effect is seen before viral protein expression. **(A)**. N6 cells were stimulated in bulk with vorinostat (3uM), prostratin (3uM) or TNFα (100ng/mL) for 24 hr. before staining for the virally encoded HSA marker protein, and flow sorting for single cells. Each cell was then assayed by qPCR for viral RNA (vRNA). Y axis values represent the fluorescent intensity of HSA staining for each cell. Each dot represents a single cell. The lower limit of quantification for vRNA is shown by a vertical dashed line. **(B)**. The percent vRNA+ cells was calculated from the percentage of cells that exhibited vRNA amplification, while percent HSA+ was calculated by gating for HSA+ cells using FlowJo analysis software. 144 cells were analyzed for each condition.

### Viral RNA and protein expression increases in an exposure time - and concentration-dependent manner

To understand vRNA expression during latency reversal at the single cell level in more detail, we examined the response of N6 cells to vorinostat at different times after stimulation (Figure 2A) and at different concentrations (Figure 2B). From baseline to 6hr. post stimulation, little change occurred in the percentage of cells expressing detectable vRNA. At 12 hr., a significant subset (40%) of cells began to express vRNA, even though HSA expression levels had not yet markedly increased.

**Fig 2.**
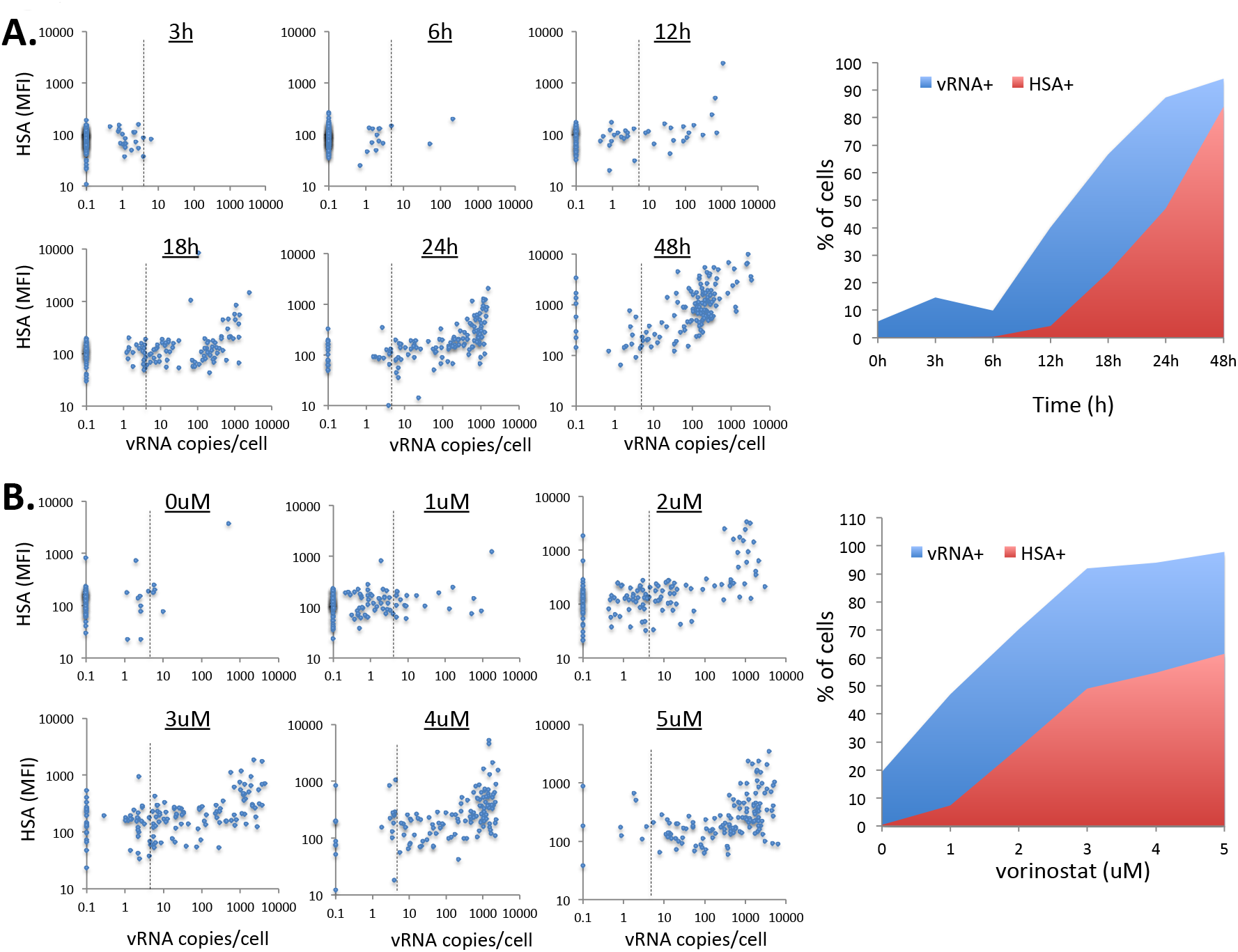
Viral RNA and protein expression increases in an exposure time - and concentration-dependent manner. **(A)**. N6 cells were stimulated with 3 μM vorinostat and, at times indicated, stained for HSA expression. Individual cells were then flow sorted and analyzed by qPCR for vRNA. **(B)**. N6 cells were stimulated with the concentrations of vorinostat indicated. At 24 hr. the cells were stained for HSA and individual cells flow sorted for qPCR of vRNA. The percent vRNA+ cells at each timepoint/concentration was calculated from the percent of cells that exhibited vRNA amplification, while percent HSA+ was calculated by gating for HSA+ cells using FlowJo analysis software. 144 cells were analyzed for each timepoint or concentration. The lower limit of quantification for vRNA is shown by a dashed vertical line.

By 18 hr., however, the proportion of vRNA+ cells had increased to 70%, and HSA expression was clearly detectable in cells expressing >500 copies of vRNA. Expression of vRNA continued to increase at 24 hr., by which time 90% of the cells were vRNA+ and ~50% were HSA+. By 48 hr., the majority of cells were both vRNA+ and HSA+, although vRNA expression levels per cell dropped significantly for HSA+ cells at this time compared to 24 hr. This observation is consistent with the findings of Martrus and coworkers (Martrus et al., 2016) and could reflect depletion of vRNA from infected cells by vRNA packaging and virion release, or reestablishment of latency in these cells. These data demonstrate significant variation in the kinetics of reactivation of vRNA expression among individual cells - vRNA levels increase rapidly (6 hr-12 hr) in a subset of cells, but slowly in other subsets of cells (24 hr-48 hr). Furthermore, exposure of N6 cells to a range of vorinostat concentrations revealed that HSA expression underestimated the percent vRNA responding cells at all concentrations (Figure 2B). For example, a significant fraction of cells (47%) expressed vRNA in response to 1 μM vorinostat, even though HSA expression at this concentration was minimal. Overall, these findings indicate that in an unstimulated latently infected cell line, vRNA expression at a low level (less than 10 copies/cell) can be detected in a small population of cells. Following LRA exposure, a larger population of cells can respond, producing viral RNA in a time- and concentration-dependent manner, and that this response precedes that which can be detected by protein-based measurements of latency reversal.

### Establishment of a primary CD4+ T cell model for HIV latency

To further examine the behavior of latently infected cells we established a primary CD4+ T cell model of HIV latency. This model is similar to models from other laboratories (Kim et al., 2014; Mohammadi et al., 2014; Sahu et al., 2006; Tyagi et al., 2010), and involves infecting activated CD4+ T cells with an eGFP expressing HIV strain (Yang et al., 2009) and sorting to obtain a pure infected population, followed by long-term (8-12 weeks) co-culture with H80 cells (Figure 3A). During this period of culture a latently infected (GFP-) population emerges. Interestingly, we observed highly variegated silencing of HIV gene expression within the infected population. Some cells exhibited complete silencing of GFP expression (GFP-), while others maintained intermediate or high levels of expression (Figure 3B, 3C). To test the ability of the latently infected cells to reactivate viral gene expression in response to T cell receptor (TCR) engagement, we purified GFP-cells from the infected cell culture and stimulated them with anti-CD3/CD28 beads. These cells reexpressed GFP to nearly 90% by 3 days, indicating that they were indeed latently infected, and that initial loss of GFP expression was not due to deletion of the provirus (Figure 3D).

**Fig 3.**
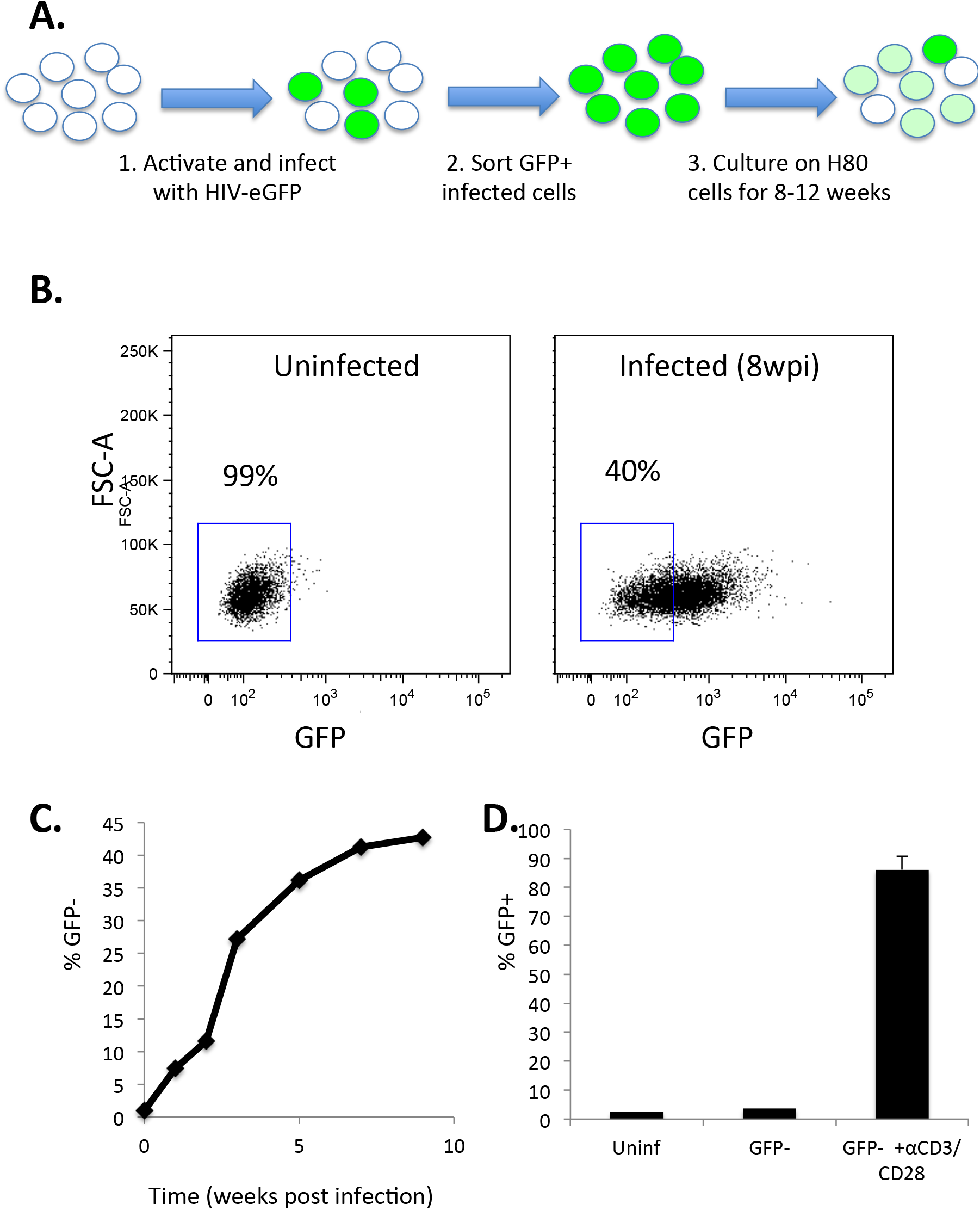
Primary CD4 T cell model of HIV latency. **(A)**. Graphical depiction of primary cell model of latency. Cells were infected with pNL4-3-Δ6-dreGFP, an eGFP expressing HIV clone, and infected (GFP+^)^ cells isolated by flow sorting. **(B)**. After 8-12 weeks of co-culturing the sorted GFP+ population with H80 cells, cells displayed heterogeneous levels of virally encoded GFP expression, with 40% now being GFP-. **(C)**. The percent GFP-cells over time was determined by flow cytometry. Data shown is a representative sample from one of three separate donors. **(D)** Latently infected cells were isolated by flow sorting of GFP-cells from the infected cell culture. These cells were then stimulated with anti-CD3/CD28 beads for 3 days. GFP expression was measured by flow cytometry before (GFP-) and after stimulation (GFP-+ αCD3/CD28), and compared to uninfected cells (uninf).

### Single cell analysis of vRNA in latently infected primary CD4+ T cells

To investigate the diversity of vRNA expression in latently infected primary cells, we analyzed sorted single cells from the infected population at 12 weeks post-infection (wpi; Figure 4). For comparison, we also examined productively infected cells at 2 days post-infection (dpi). We found vRNA levels in primary CD4+ T cells were linearly correlated with GFP protein expression for both timepoints (r^2^=0.179, p=1.71×10^−7^ t-test for 12 wpi, r^2^=0.475, p=1.35×10^−21^ for 2dpi). This contrasts with the apparent threshold of vRNA required for viral protein expression in N6 Jurkat cells. The 2 dpi population expressed higher overall levels of vRNA than cells at 12wpi (p <0.00001, Mann-Whitney test) indicating that transcriptional downregulation accounted for part of the reduction in GFP protein expression over time, but interestingly, a subset of 12 wpi cells expressed vRNA to a similar level to that seen at 2 dpi. At 12 wpi, 12% of cells exhibited undetectable vRNA, indicating transcriptional silencing. This analysis shows that after long periods in culture, most infected cells downregulate viral gene expression at both the RNA and protein level but the extent of this silencing varied greatly between individual cells. This observation indicates considerable heterogeneity in the expression level of vRNA within a latently infected primary cell culture

**Fig 4.**
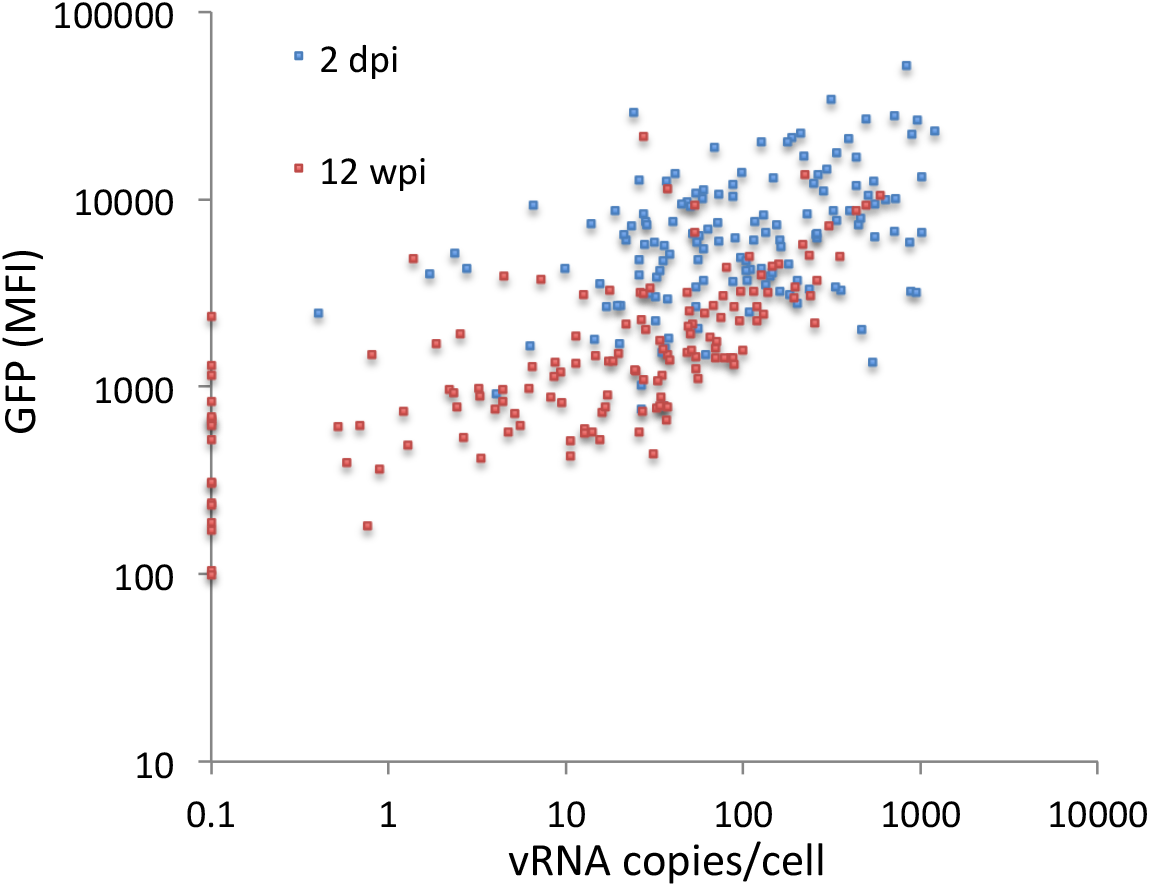
Single cell viral RNA and antigen expression in the primary cell latency model is heterogeneous. Single infected primary CD4 T cells at 2 dpi or 12 wpi were flow sorted and analyzed for vRNA expression by sc-qPCR. GFP fluorescent intensity (MFI) for each cell was plotted against vRNA copies per cell. Each dot represents data from an individual cell.

### Differential proviral transcription in cells with different host cell gene expression patterns

We hypothesized that the heterogeneity of proviral expression that we observed in the primary cell latency model was related to underlying heterogeneity within the primary cells themselves. CD4+ T cells are a diverse collection of cells, with different subsets, developmental stages and sub-states that could impact HIV transcription. Additionally, stochastic biological noise and random fluctuation in the expression of CD4+ T cell transcription factors could also potentially affect HIV expression. To test this hypothesis, we performed single-cell RNA-Seq (scRNA-Seq) on 2,676 infected cells and 3,097 control uninfected cells at 12 weeks post infection. Within the infected population, approximately 38.2% of cells were GFP- by flow cytometry, while the remaining 61.8% expressed varying levels of GFP.

We first compared the single-cell transcriptomes of the uninfected and infected cell populations, after 12 weeks in culture. Unsupervised graph-based clustering and reduction of the dimensionality of the transcriptomes of single-cells to two dimensions by t-distributed Stochastic Neighbor Embedding (tSNE) created a single-cell map of clusters of cells. Overall, the infected cells were highly similar to the uninfected cells with regard to transcriptome expression, and clustered together (Figure 5A). Out of 21,258 total genes detected in both samples, we identified 14 upregulated and 15 downregulated transcripts when comparing infected and uninfected cells (Figure 5B, Table 1). As a marker of viral expression, GFP was the most differentially expressed transcript followed by *IFITM1* and *MIF* (Figure 5C). *IFITM1* has been shown to be upregulated upon HIV infection and even identified as a candidate marker of latently infected cells (Raposo et al., 2017) and *MIF* has been shown to be elevated in plasma from HIV-infected individuals and may play a role in viral replication (Regis et al., 2010). By contrast, antiviral molecules *GNLY* and *ISG15* and ribosomal transcript *RPL36A* were expressed at lower levels in infected cells (Figure 5C).

**Fig 5.**
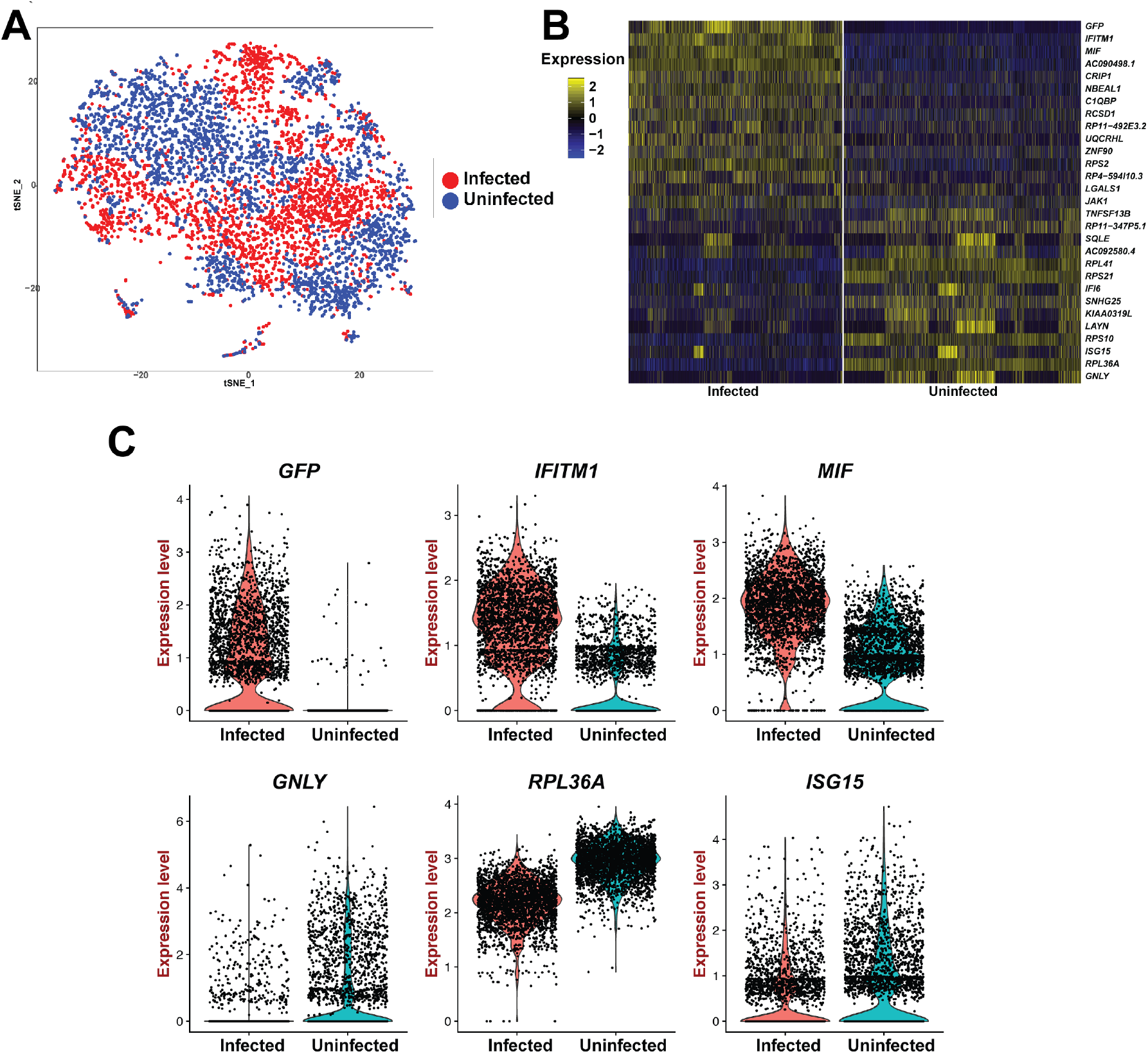
Latently infected cells exhibit modest changes compared to uninfected cells. **(A)**. Two-dimensional plot from unsupervised clustering by t-distributed stochastic neighbor embedding (tSNE) of the single-cell transcriptomes from 2,676 infected (red) and 3,097 uninfected (blue) primary CD4 T cells. **(B)**. Heatmap of significantly changed transcripts between infected and uninfected cells (likelihood ratio test; P ≤ 0.05). Each pixel column is the expression of an individual cell. Transcripts are on the rows with the normalized expression (Z-scores) colored by the legend (yellow, more upregulated; blue, more downregulated). **(C)**. Violin plots detailing the expression levels of the top 3 upregulated and downregulated transcripts within infected cells.

**Table 1:** Transcripts differentially expressed in infected vs uninfected. The set of genes that are differentially expressed between infected and uninfected cells was determined. Genes are ranked by average log fold change. p values were calculated by a Likelihood ratio test.

| Gene | p value | average log difference |
| --- | --- | --- |
| GFP | 0 | 1.370931972 |
| IFITM1 | 0 | 0.987611872 |
| MIF | 0 | 0.942373398 |
| AC090498.1 | 0 | 0.908621278 |
| CRIP1 | 3.09E-242 | 0.503730152 |
| NBEAL1 | 9.22E-221 | 0.438055216 |
| C1QBP | 2.88E-150 | 0.344013897 |
| RCSD1 | 6.88E-127 | 0.343583613 |
| RP11-492E3.2 | 6.89E-60 | 0.320113293 |
| UQCRHL | 2.09E-142 | 0.287809102 |
| ZNF90 | 4.48E-98 | 0.285478496 |
| RPS2 | 4.19E-308 | 0.271615274 |
| RP4-594I10.3 | 5.58E-96 | 0.263319428 |
| LGALS1 | 1.03E-23 | 0.260550424 |
| JAK1 | 6.47E-76 | 0.256058832 |
| TNFSF13B | 6.72E-39 | -0.261485221 |
| RP11-347P5.1 | 3.78E-78 | -0.270064479 |
| SQLE | 9.64E-30 | -0.295511823 |
| AC092580.4 | 3.85E-47 | -0.304995107 |
| RPL41 | 0 | -0.309907065 |
| RPS21 | 0 | -0.346851961 |
| IFI6 | 1.90E-41 | -0.372868853 |
| SNHG25 | 1.38E-146 | -0.382496193 |
| KIAA0319L | 8.45E-90 | -0.388877371 |
| LAYN | 1.52E-60 | -0.426304333 |
| RPS10 | 0 | -0.452350093 |
| ISG15 | 1.86E-50 | -0.470881381 |
| RPL36A | 0 | -0.74141456 |
| GNLY | 3.41E-155 | -1.348770687 |

Next, we focused our analysis on the heterogeneity within the infected cell population. Using graph-based clustering, infected CD4+ T cells formed 5 distinct transcriptional clusters (Figure 6A). Strikingly, we observed that viral gene expression, as measured by *GFP* RNA levels, was significantly different between the clusters. While we identified cells exhibiting proviral expression in all 5 clusters, we found that cluster 1 had significantly (*P* < 0.001; likelihood-ratio test) higher expression of viral *GFP* compared to the other four clusters and the lowest expression within cluster 0 (Figure 6B & 6C). These data suggested that viral gene expression was impacted by the host cell transcriptomic profile. Next, we identified transcripts that were significantly enriched within each cluster (Figure 6D, Supplemental table 1). Over the 5 clusters, 1213 genes were differentially expressed between individual clusters. Cluster 0 was defined by a signature of 39 genes, while cluster 1 was defined by a signature of 141 genes (Supplemental Table 1). Notably, cluster 0, with the lowest level of HIV vRNA expression, contained cells expressing high levels of known markers of T cell lymph node homing (CCR7, CXCR4, and SELL (CD62L)), as well as the cytokine receptor CD127. These genes are known to be expressed to high levels in naïve and central memory T cells, suggesting that cluster 0 is enriched with cells with a naïve or Tcm phenotype (Figure 6D). By contrast, cluster 1 with the highest level of vRNA, contained several markers of T cell activation, such as *HLA-DR, IL2RA* (CD25) *TNFRSF4* (0X40L) and *TNFRSF18* (GITR) (Figure 6D, Supplemental table 1). These data suggested a relationship between the level of HIV transcriptional activity in the infected cell and the transcriptome of the infected host cell.

**Fig 6.**
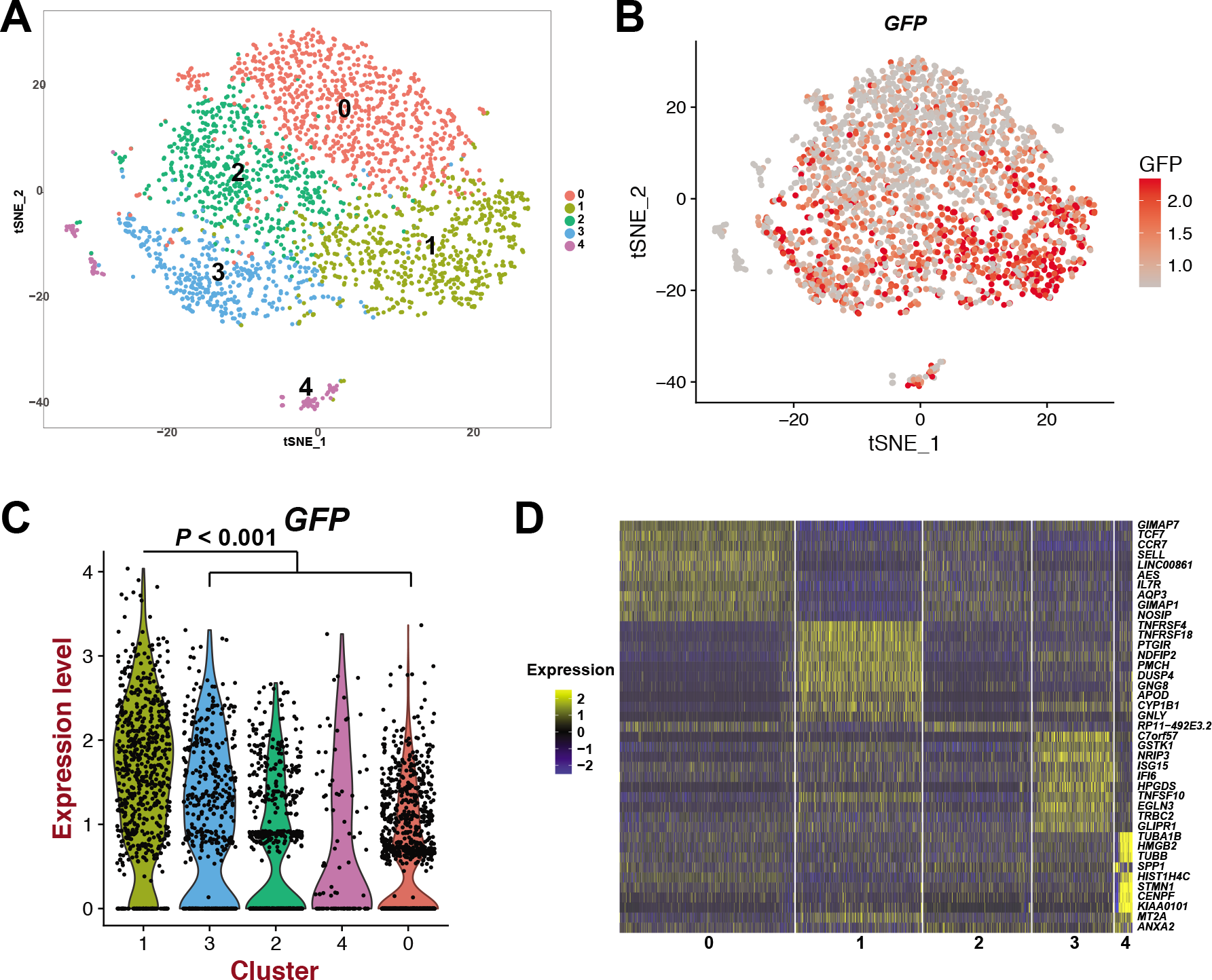
Differential proviral expression levels in clusters with different transcriptomic profiles. **(A)**. Graph-based clustering of 2,676 infected cells represented on the tSNE plot. **(B)**. Normalized transcript expression level of *GFP* on the tSNE plot representation. Legends indicate normalized transcript values. **(C)**. Violin plots of *GFP* expression within the identified clusters. **(D)**. Heatmap of the top 10 transcripts that define each cluster. Panels represent individual clusters and each pixel column is the expression of individual cells. Transcripts are on the rows with normalized expression (Z-scores) colored by the legend (yellow, more upregulated; blue, more downregulated).

### HIV transcription is preferentially silenced in cells with greater proliferative potential

Next, to further describe the association of HIV gene expression with host cell gene expression, we identified transcripts that were differentially expressed between cells expressing detectable levels of vRNA (GFP+) or undetectable levels of vRNA (GFP-), independent of their prior assignment to Cluster 0 through 5 above. We identified 34 transcripts upregulated and 13 transcripts downregulated in GFP+ cells compared to GFP-cells (Table 2). Consistent with the cluster analysis described above, GFP-cells had higher expression of markers associated with naïve or central memory T cell phenotype such as *CCR7* and *SELL* and lower expression of markers of activated T cells such as *IL2RA, HLA-DR, CD38* and *CD96* (Figure 7A & 7B). To investigate whether the set of genes that differentiate transcriptionally silent HIV infection (GFP-) from transcriptionally active infection (GFP+) represented differential activity of specific biological pathways, we preformed Ingenuity Pathway analysis (IPA) (Jiménez-Marín et al., 2009). Strikingly, this analysis demonstrated significant enrichment (p<10^−8^) for expression of genes that contribute to cell survival and proliferation in cells with lower HIV transcription (Figure 8A, Supplemental Figure 1). Based on this analysis we hypothesized that HIV transcriptional silencing might be correlated with differential survival or proliferative potential of the infected cells. To test this hypothesis, we sorted infected cells at 8 wpi into two populations based on GFP expression level (GFP-, GFP-low and GFP-high) (Fig 8B; **left panel**). We then stimulated each population with anti-CD3/CD28 beads in the presence of IL-2 100 U/mL for three days, then allowed the culture to expand further in the absence of stimulation. After 14 days of expansion, we measured expansion by counting viable cells. Notably, GFP-cells exhibited the greater fold expansion of the two cultures (Figure 8B, right panel). These data confirmed that HIV transcriptional silencing occurs preferentially in a specific subset of cells with greater proliferative potential, and are consistent with the hypothesis that HIV transcriptional silencing in primary CD4+ T cells is associated with intrinsic biological properties of the host cells.

**Fig 7.**
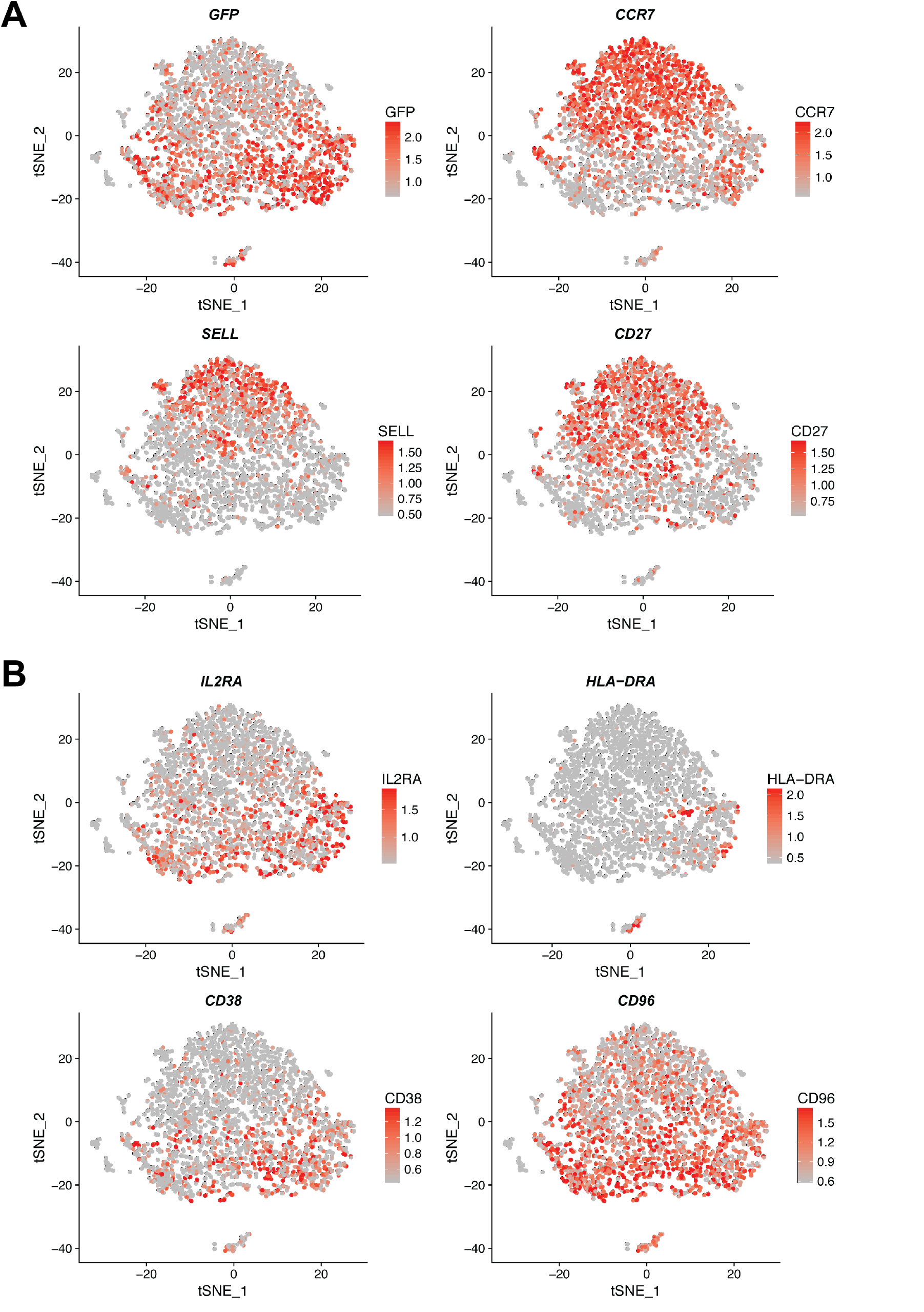
Expression patterns of host cell genes that associate with HIV transcription level. Normalized expression pattern for selected genes whose expression negatively **(A)**, or positively **(B)**, correlated with viral gene expression (GFP) plotted on the tSNE plot representation. Each dot represents and individual cell. Legends indicate normalized transcript values.

**Fig 8.**
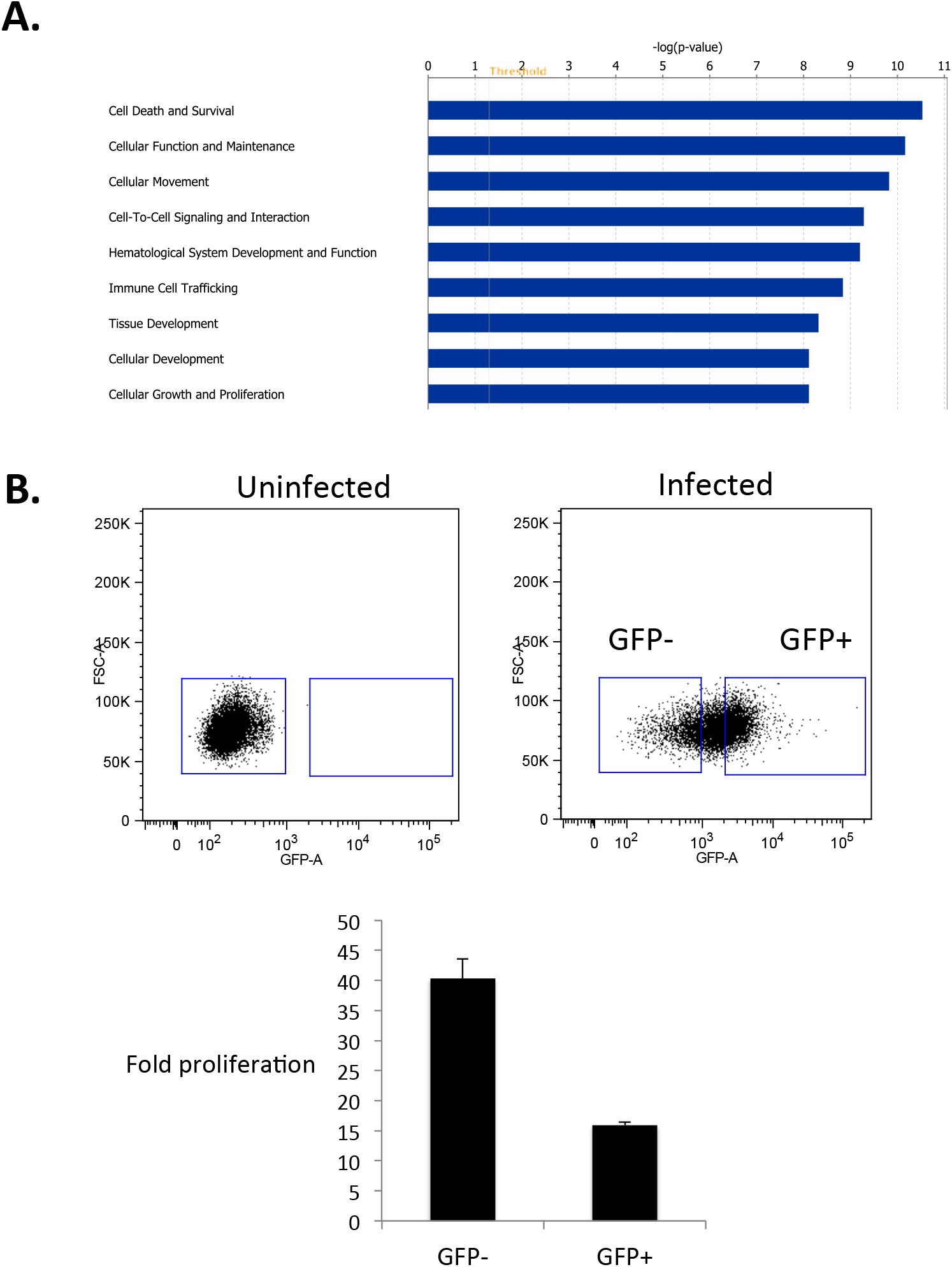
HIV is preferentially silenced in cells with higher proliferative capacity. **(A)**. To identify cellular pathways/functions that were enriched in genes associated with HIV silencing, Ingenuity Pathway analysis (IPA) was used to analyze the set of genes that are differentially expressed between cells with active viral gene expression (GFP+) and those with silenced viral gene expression (GFP-), and the most prominently affected cellular functions are shown. **(B)**. An infected culture of primary CD4 T cells at 8wpi was sorted into two populations based on the level of GFP expression. - GFP - and GFP+ (upper right panel). Equivalent numbers of sorted cells were then stimulated with αCD3/αCD28 beads for three days and then expanded in 100 U/mL IL-2 for two weeks. At the end of this period the fold expansion for each culture was calculated (lower panel). Data shown are representative of three independent donors.

**Table 2:** Transcripts differentially expressed between GFP+ and GFP-infected cells. The set of genes that are differentially expressed between GFP+ and GFP-cells within the infected cell population was determined. Genes are ranked by average log fold change. p values were calculated by a Likelihood ratio test.

| Gene | p value | average log difference |
| --- | --- | --- |
| GFP | 0 | 1.699930123 |
| PMCH | 2.44E-28 | 0.678372138 |
| TNFRSF18 | 1.73E-39 | 0.536725908 |
| KLRB1 | 5.74E-19 | 0.492116133 |
| TNFRSF4 | 1.69E-37 | 0.470618182 |
| AC092580.4 | 9.12E-38 | 0.459071737 |
| ALOX5AP | 1.76E-35 | 0.451496403 |
| CYP1B1 | 1.25E-24 | 0.421925717 |
| SRGN | 2.13E-48 | 0.403167028 |
| NDFIP2 | 9.86E-41 | 0.389106789 |
| PTGIR | 3.76E-30 | 0.368909492 |
| DUSP4 | 2.51E-28 | 0.361458251 |
| IL2RA | 1.31E-23 | 0.354081966 |
| GAPDH | 8.16E-32 | 0.343675603 |
| ENTPD1 | 6.22E-33 | 0.339960123 |
| GNG8 | 3.76E-22 | 0.333732314 |
| NR3C1 | 5.43E-30 | 0.331914346 |
| ID2 | 4.19E-25 | 0.319738595 |
| IL32 | 6.77E-37 | 0.314489333 |
| TNFSF10 | 2.13E-22 | 0.308558414 |
| AHI1 | 7.99E-21 | 0.30770296 |
| TMSB10 | 3.10E-31 | 0.299907141 |
| BIRC3 | 1.05E-15 | 0.294641539 |
| AIF1 | 2.11E-24 | 0.289685315 |
| TRBC1 | 2.99E-12 | 0.287790466 |
| CTSH | 1.12E-22 | 0.274799529 |
| HLA-DPA1 | 8.80E-27 | 0.274508816 |
| PPFIBP1 | 4.36E-24 | 0.26652867 |
| SMCO4 | 3.33E-24 | 0.266433234 |
| LINC00152 | 2.85E-22 | 0.260834793 |
| PDE7B | 5.33E-25 | 0.258575644 |
| CLIC1 | 2.84E-23 | 0.253523063 |
| UNQ6494 | 2.16E-29 | 0.253168389 |
| CD9 | 4.23E-12 | 0.253001841 |
| LTB | 8.92E-17 | -0.264957724 |
| GIMAP7 | 3.50E-23 | -0.269112444 |
| PLP2 | 9.53E-29 | -0.269151701 |
| SELL | 5.51E-18 | -0.273430759 |
| AES | 1.60E-26 | -0.275624352 |
| S100A10 | 6.70E-23 | -0.280085029 |
| NGFRAP1 | 7.14E-23 | -0.290697668 |
| MT-ND2 | 1.99E-36 | -0.292218933 |
| OCIAD2 | 1.71E-42 | -0.292294496 |
| CCR7 | 1.86E-20 | -0.297581957 |
| ANXA2 | 2.39E-39 | -0.416759518 |
| SPP1 | 7.96E-48 | -0.568447373 |
| S100A6 | 8.99E-73 | -0.721523376 |

### Viral silencing is associated with T cell subset identity

The finding that gene expression can define specific T cell subsets within a cluster of infected cells with the lowest level of HIV gene expression suggested a relationship between specific CD4+ T cell subsets and silencing of the HIV promoter. CD4+ T cells emerge from the thymus as naïve T cells (Tn), then, through a combination of antigenic simulation and cytokine cues, develop linearly into central memory T cells (Tcm), transitional memory T cells (Ttm), and finally effector memory T cells (Tem). These subsets are defined by differential expression of a set of surface markers including CD45RO, and CCR7. Unfortunately, our scRNAseq data was unable to accurately distinguish between the naïve cell marker CD45RA and the memory cell marker CD45RO due to these alternatively spliced transcripts possessing identical 3’ sequences. Thus, to examine this hypothesis, we examined expression of CCR7 and CD45RO on the surface of infected cells by flow cytometry after 8 weeks of coculture with H80 cells. This staining confirmed that the infected cell population consisted of a mixture of cells with “naïve” (CCR7+ CD45RO-) phenotype, cells with a Tcm phenotype (CCR7+ CD45RO+), and cells with a Tem/Ttm (CCR7-, CD45RO+) phenotype (Figure 9, left panel). Notably, the distribution of cells within these populations differs from other reports using a similar model (Tyagi et al., 2010; Yang et al., 2009). This difference is likely explained by methodological differences in the preparation of the cells - in one of the previous studies (Tyagi et al., 2010), the infected cells were reactivated and expanded post sorting, while in our experiments, the infected cells were not re-stimulated after the initial activation. To determine the level of viral gene expression in each subset, we examined the percent of GFP-(latent) cells in each subset (Figure 9, right panel). All three populations exhibited a mixture of GFP- and GFP+ cells, but, consistent with scRNAseq, we observed that there was a gradient of HIV silencing across the populations, with CD45RO- CCR7+ cells having the highest percentage of GFP-cells, and CD45RO+ CCR7-cells exhibiting the lowest. These results demonstrated that, in this latency model, HIV transcription was preferentially silenced in cells at the Tn/Tcm end of the developmental spectrum. Nevertheless, we also observe heterogeneous viral expression levels within each subset, consistent with the hypothesis that latency is the output of a convergence of factors at the level of each individual cell. In this model, we would expect a diverse population of latently infected cells, the most frequent phenotype of which would reflect the most common convergence of these influences: i) a cell that had recently returned to the central memory pool from a highly activated effector population prone to the initial stages of HIV infection, and ii) a provirus whose expression had become restricted by epigenetic marks.

**Fig 9.**
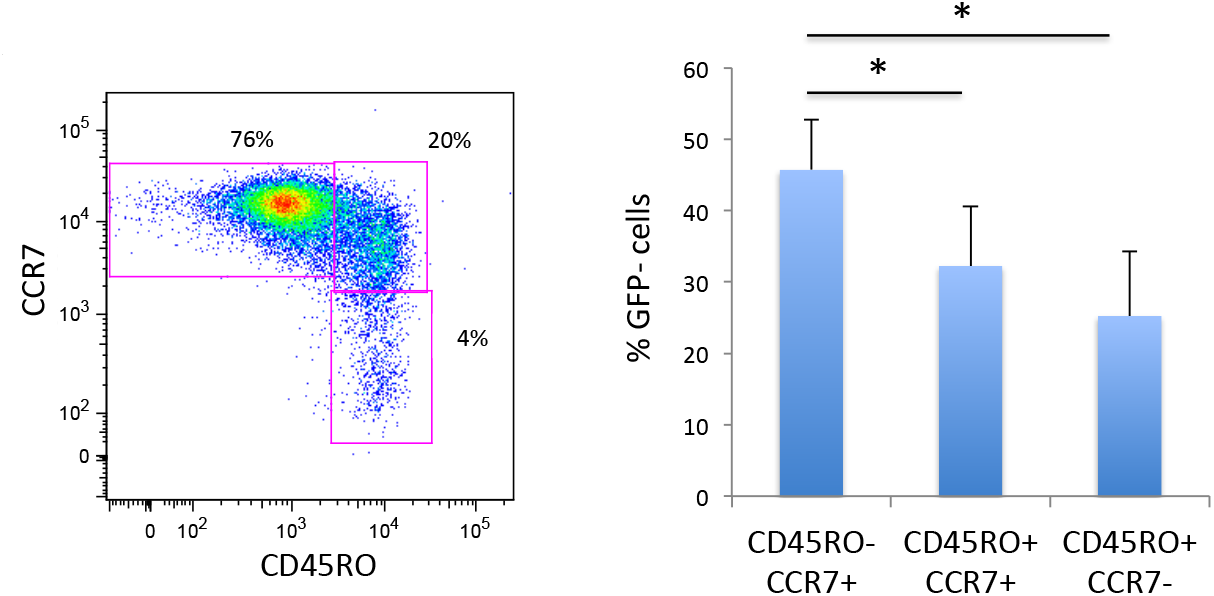
HIV is preferentially silenced in cells expressing Tn and Tcm markers. Infected cells at 8wpi were stained for surface markers CD4, CD45RO and CCR7 to identify different CD4 T cell subsets. CD45RO-, CCR7+ cells represent naïve T cells (Tn), CD45RO+, CCR7+ cells represent central memory cells (Tcm), while CD45RO+, CCR7-cells represent a mix of transitional memory T cells (Ttm) and effector memory T cells (Tem). The fraction of GFP-cells in each gate was then calculated and plotted (right panel). Data shown in the right panel represent the average of four independent donors. An asterisk represents a p value of <0.05, Students T-test.

## Discussion

In this study we have probed the expression of viral and host cell genes in individual latently infected cells. These data reveal highly heterogeneous transcription of the provirus within the population, and that this heterogeneity is associated with distinct host cell transcriptional signatures. Over the past few years, single cell methods have become widely used, and have provided powerful new insights into heterogeneity in biological systems that were previously assumed to be homogeneous (Linnarsson and Teichmann, 2016; Shalek et al., 2014; Villani and Shekhar, 2017). These methods are particularly useful to the study of virus-host interactions, where multiple layers of complexity arise from variation in both host cells and infecting viruses (Ciuffi et al., 2016). The HIV latent reservoir, for example, consists of thousands of genetically distinct viral genomes, including those encoding immune escape variants, that persist as transcriptionally silent proviruses integrated into a range of genomic locations. Furthermore, expression of these proviruses that are regulated by a wide range of dynamic chromatin modifications and host cell transcription factors that can be divergently expressed and differentially active in different CD4+ T cell subpopulations. The latent HIV reservoir thus represents not a single, uniform target, but a diverse mixture of several subpopulations of infected cells, potentially with fundamentally different characteristics. The development of strategies to reactivate or eliminate these cells may need to account for this diversity within the latent reservoir. For example, targeting this reservoir with latency reversing agents may require separate strategies for different subtypes of cells rather than a “one size fits all” approach. Consistent with this notion, upon a single round of induction, individual LRAs typically disrupt latency in only a fraction of replication competent proviruses (Ho et al., 2013). As such there is an urgent need to study latency in the context of approaches that can observe and characterize individual latently infected cells.

Prior publications have also applied single cell approaches to the study of HIV latency, although with significantly different methodologies. Weigand et al 2017 used cell-associated HIV RNA/DNA quantitation combined with single HIV genome sequencing, both at limiting dilution, to detect diversity of viral transcription from individual proviruses within samples from patients on ART (Wiegand et al., 2017). Similar to our results, they found that a minor subset of latently infected cells make detectable RNA in the absence of stimulation and that vRNA levels different greatly from infected cell to infected cell. Likewise, Bui et al 2015 found that burst size of viruses release from single infected cells varies over a wide range (Bui et al., 2015). Flow cytometry using probes for viral transcripts has also yielded insight into diversity of virus and host gene expression in individual cells during infection (Baxter et al.; Bolton et al., 2017; Martrus et al., 2016).

Using a single cell vRNA assay, we find that viral transcription levels are highly heterogeneous between individual latently infected cells. This observation was true for both a Jurkat derived cell line (N6) as well as latently infected primary cells. In both model systems, a sizeable subpopulation of latently infected cells transcribed low levels of viral RNA in the absence of stimulation. Furthermore, the data demonstrate that antigen-based assays for latency reversal significantly underestimate the fraction of responding cells after stimulation with LRAs. N6 cells exhibited an apparent threshold of vRNA expression before virally encoded antigen became detectable, while for primary cells, this relationship was more complex. The reason for this difference is unclear, but could be related to the greater levels of underlying heterogeneity in primary cells.

In our primary cell latency model, we observed only minor differences between infected and uninfected cells using scRNAseq, suggesting that viral reprogramming of infected cells during latency is limited. This finding is consistent with a previous population analysis of the transcriptome of infected cells using a similar model (Mohammadi et al., 2014). However, the virus used in these studies lacks expression of most viral proteins in order to limit cytopathic effect, and thus may not reflect the full impact of an intact replication competent virus on the host cell. By contrast, we observed a striking association between activity of the HIV promoter and the host cell transcriptome within the infected cell population. Thus, our data argue that at least part of the basis for heterogeneous viral transcription during latency is explained by underlying heterogeneity within the infected host cells. In particular, we observed a clear preference for HIV silencing in cells expressing markers of Tn and Tcm subsets, while effector and activated cells were associated with higher levels of viral gene expression. These results suggest a role for T cell subset identity and intracellular environment in regulating the outcome of infection. Consistent with this notion, it has recently been demonstrated that HIV preferentially enters latency if infection occurs during a period of global cellular transcriptional downregulation as cells return to rest from activation (Shan et al., 2017). It is unlikely that the preferential silencing we observe in Tn and Tcm in our primary cell model is related to differential mutagenesis of the provirus, since almost 90% of sorted latent (GFP-) primary cells in our model re-expressed GFP upon TCR stimulation. Our finding of preferential silencing in Tn and Tcm cells is also consistent with outgrowth studies from suppressed patents (Soriano-Sarabia et al., 2014).

The molecular basis for differential silencing across CD4+ T cell subtypes is unclear. As T cells develop along a linear developmental trajectory from Tn to Tcm, then Ttm and Tem cells, they undergo progressive epigenetic remodeling, characterized by de-repression of cellular genes and loss of histone methylation islands that regulate expression of these genes (Durek et al., 2016). Thus, the pattern of HIV silencing that we observe mirrors with the overall epigenetic program of these cells, suggesting a link between the two. In addition to epigenetic differences, T cell subsets may also exhibit differences in the behavior of other important mechanisms of HIV transcriptional regulation, such as Tat, or transcriptional elongation (Kim et al., 2011; Razooky et al., 2015; Weinberger et al., 2005; Yukl et al., 2018). Nevertheless, with each subset, we see a mixture of GFP+ and GFP-cells, indicating that subset identity may make a given provirus vulnerable to viral silencing, but that other factors must also contribute. The functional role of individual genes whose expression is enriched in the GFP-and GFP+ clusters in HIV gene expression should be investigated. Notably, we also observe transcriptional heterogeneity within N6 cells, a clonal population of Jurkat cells. Specifically, a subpopulation of these cells transcribes viral RNA in the absence of stimulation, and they also exhibit widely different vRNA levels after stimulation. Furthermore, we observe that these cells can demonstrate heterogeneous kinetics of response to LRA stimulation. These cells likely possess underlying heterogeneity with respect to host cell gene expression, due to asynchronous cell cycling, or from random biological noise. Given that all N6 cells eventually reactivated and expressed viral antigen, it is probable that part the diversity of vRNA expression by these cells arises from variegated delays to reactivation.

An important caveat to these and all such observations, is that they are generally made over limited periods of time, after a single exposure to LRA. As LRAs are now being testing in the clinic in cyclical, repeated exposures over weeks, investigations will have to come to understand the cumulative effects of LRA exposure over time. And if the studies *in vivo* also reveal that transcriptional responses are more frequent and diverse than can be appreciated by the detection of a reporter gene product or a viral antigen, then the relevant threshold of latency reversal in vivo will not be that at which viral transcription can be measured, but that at which the immune system can detect and respond to viral antigen expression over time. These insights give hope that, while complex, a deeper understanding of the biology of HIV latency may lead to effective therapeutic approaches to clear persistent proviral infection.

## Acknowledgements

Research reported in this publication was supported, in part, by CARE, a Martin Delaney Collaboratory of the National Institute of Allergy and Infectious Diseases (NIAID), National Institute of Neurological Disorders and Stroke (NINDS), National Institute on Drug Abuse (NIDA) and the National Institute of Mental Health (NIMH) of the National Institutes of Health, grant number 1UM1AI126619-01. The content is solely the responsibility of the authors and does not necessarily represent the official views of the National Institutes of Health. We thank Nancy Fisher and the UNC Flow Cytometry Core Facility, supported in part by P30 CA016086 Cancer Center Core Support Grant to the UNC Lineberger Comprehensive Cancer Center, and supported in part by the North Carolina Biotech Center Institutional Support Grant 2012-IDG-1006. This work was also supported by the UNC Center for AIDS Research (P30 AI50410). We acknowledge Connor Hart and Bhavna Hora for technical assistance and the Duke Human Vaccine Institute Viral Genetic Analysis Core.

## Methods

### Single cell viral RNA (sc-vRNA) assay

Cells were stained at 1:1000 in phosphate buffered saline (PBS) with the live/dead dye Zombie Violet (ZV; Biolegend, San Diego CA) for 20mins. Cells were then washed and resuspended in staining buffer (PBS with 2% fetal calf serum and 1mM EDTA). Next, single cells were sorted into 96-well PCR plates with 10 μl of TCL buffer (Qiagen, Hilden, Germany) containing 1% Beta-mercaptoethanol using an Aria flow sorter (Becton Dickinson, Franklin Lakes, NJ). Doublets were identified and excluded by forward scatter (FSC)/side scatter (SSC) gating, and dead cells were excluded by gating on ZV-cells. Index sorting was used to record fluorophore/GFP intensity for each cell. After sorting plates were briefly spun and frozen at −80°C. To extract RNA, plates were thawed and 22 μl of RNA-Clean XP beads (Beckman Coulter, Brea, CA) added and incubated for 10 min. Beads were then isolated and washed in 80% ethanol, then eluted into 10μl of 1x PCR master mix using Fastvirus (Thermo, Waltham, MA) and primer sets for HIV Gag (GAG-F: ATCAAGCAGCCATGCAAATGTT, GAG-R: CTGAAGGGTACTAGTAGTTCCTGCTATGTC, GAG-Probe: FAM/ZEN-ACCATCAATGAGGAAGCTGCAGAATGGGA-IBFQ) and Betaactin (BAC-F: TCACCCACACTGTGCCCATCTACGA, BAC-R: CAGCGGAACCGCTCATTGCCAATGG, BAC-Probe: HEX-ATGCCCTCCCCCATGCCATCCTGCGT-IBFQ). This plate was then run on a QS3 (Applied Biosystems, Foster City, CA) real time thermocycler with a 5 minute reverse transcription step at 50°C, followed by 40 cycles of 94°C (3 sec.), 60°C (30 sec.). RNA copy number was determined by comparison to a standard curve of synthesized Gag gblock purchased from Integrated DNA Technologies (Coralville IA). Wells that failed to amplify Beta actin were excluded from analysis. The lower limit of quantification (LLOQ) for this assay was seven copies of Gag RNA. Coefficient of variation for technical replicates in the standard curve ranged from 10-15% for 5000 copies to 40-45% for 7 copies. Analysis of uninfected cells indicated a false positive amplification rate of approximately 1%, with the majority of false positive signals falling below the LLOQ.

### Cell lines and culture

N6 cells were a generous gift from David Irlbeck (Glaxo SmithKline, Chapel Hill, NC). This cell line was derived from Jurkat cells and was made using HIV-1 engineered to express a luciferase reporter in place of the HIV-1 nef gene with an additional mouse heat stable antigen CD24 (HSA) reporter located just downstream of the luciferase open reading frame, separated by a T2A element (NLCH-Luci-HSA). NLCH, the parent molecular infectious clone was kindly provided by the laboratory of Dr. Ron Swanstrom (UNC-Chapel Hill, Chapel Hill, NC) and is a modification of HIV-1 NL4-3 (GenBank U26942) where flanking sequences were removed. To generate latently infected clones, NLCH-Luci-HSA infected Jurkat cells expressing low levels of HSA were selected, limit diluted, and individual clones, including clone N6, were expanded in culture. This cell line was maintained in RPMI media with 10% Fetal calf serum (FCS) and penicillin/streptomycin. 500nM Efavirenz was added to the media to prevent spontaneous virus outgrowth. H80 cells were obtained from Darrell Bigner (Duke, Durham NC) and maintained in RPMI media with 10% FCS and penicillin/streptomycin. 293T cells for virus production were grown in Dulbecco’s modified eagle medium (DMEM) with 10% FCS and penicillin/streptomycin.

### Antibodies/flow cytometry

For flow cytometry, cells were stained in 100 μl staining buffer (PBS with 2% FCS and 1 mM EDTA). Staining was carried out for 30 min. on ice, before washing with PBS and resuspension in staining buffer. Flow cytometry was carried out on a BD Fortessa (Becton Dickson). Antibodies/fluorophores used for staining were anti-CD24-PE, anti-CD4-BV605, anti-CD45RO-BV785, anti-CCR7-APC. All antibodies were purchased from Biolegend.

### Virus production

HIV stocks were produced by transient transfection of 293T cells with pNL43-Δ6-dreGFP plasmid, as well as the packaging plasmids PAX2 and gp160 envelope, using Mirus LT1 tranfection reagent (Mirus Bio, Madison, WIU). pNL43-Δ6-dreGFP was kindly provided by Robert Siliciano (Johns Hopkins). This virus contains premature stop codons in all viral genes except tat and rev, and contains a destabilized eGFP gene in the envelope open reading frame (Yang et al., 2009). The gp160 expression plasmid was derived from the CXCR4-tropic NL4-3 strain and was kindly provided by Ronald Swanstrom (UNC Chapel Hill). Tissue culture media was replaced at 24h. At 48 hr. supernatant was harvested and spun at 2000 rpm for 5 mins to remove cell debris before filtering through a 0.45μM filter (Millipore, Burlington MA). Aliquots of virus were frozen at −80°C.

### Primary cell model of HIV latency

Total CD4 T cells were isolated from peripheral blood mononuclear cells (PBMCs) by negative selection using Easysep total CD4 T cell isolation kit (Stem Cell, Vancouver, BC). Purity was determined by staining with anti-CD4-FITC and flow cytometry and was typically ~98-99%. For infection, 20 million CD4 T cells were activated by mixing with anti-CD3/CD28 beads (Thermo Fisher) at one bead per cell for 2 days with 100U/mL IL-2. At 2d, the beads were magnetically removed, and the cells infected with pNL43-Δ6-dreGFP virus by centrifugation at 600 g for 2h at room temperature, in the presence of 4ug/mL polybrene. At 2dpi, cells were resuspended in staining buffer, and infected (GFP+) cells were isolated using a FACSAria flow sorter (Becton Dickson). The recovered GFP+ cells were then co-cultured with H80 cells (provided by Darrell Bigner, Duke University) and 20 U/mL IL-2 for up to 12 weeks. Media for the co-culture was replaced every 2-3 days. Infected cells were moved to flasks with fresh H80 cells every two weeks.

### Single cell RNAseq

Single-cell RNAseq was performed as described (Zheng et al., 2017). Briefly, cellular suspensions were loaded on a GemCode Single-Cell instrument (10X Genomics, Pleasanton, CA) to generate single-cell beads in emulsion. Single-cell RNAseq libraries were then prepared using GemCode Single Cell 3’ Gel bead and library kit (10X Genomics). Single-cell barcoded cDNA libraries were quantified by quantitative PCR (Kappa Biosystems, Wilmington, MA) and sequenced on an Illumina NextSeq 500 (San Diego CA). Read lengths were 26 bp for read 1, 8 bp i7 index, and 98 bp read 2. Cells were sequenced to greater than 50,000 reads per cell. The Cell Ranger Single Cell Software Suite was used to perform sample demultiplexing, barcode processing and single-cell 3’ gene counting. Reads were aligned to human genome release Hg38 with GFP nucleotide sequence added as an additional gene. Graph based cell clustering, dimensionality reduction and data visualization were analyzed by the Seurat R package (Satija et al., 2015). Cells that exhibited high transcript counts (>7500 total transcripts), >0.1% mitochondrial transcripts, or transcripts characterized by the H80 feeder cells were excluded from analysis. Differentially expressed transcripts were determined in the Seurat R package utilizing the Likelihood-ratio test for single cell gene expression statistical test (McDavid et al., 2013). Single cell RNAseq data shown in the main text are representative from one of three independently analyzed donors. Additional donor data shown in supplemental information. scRNAseq data has been deposited in the Biosample database. Accession numbers are: SAMN08685499, SAMN08685500, SAMN08685501, SAMN08685502

---

**Supplemental Figure 1.**
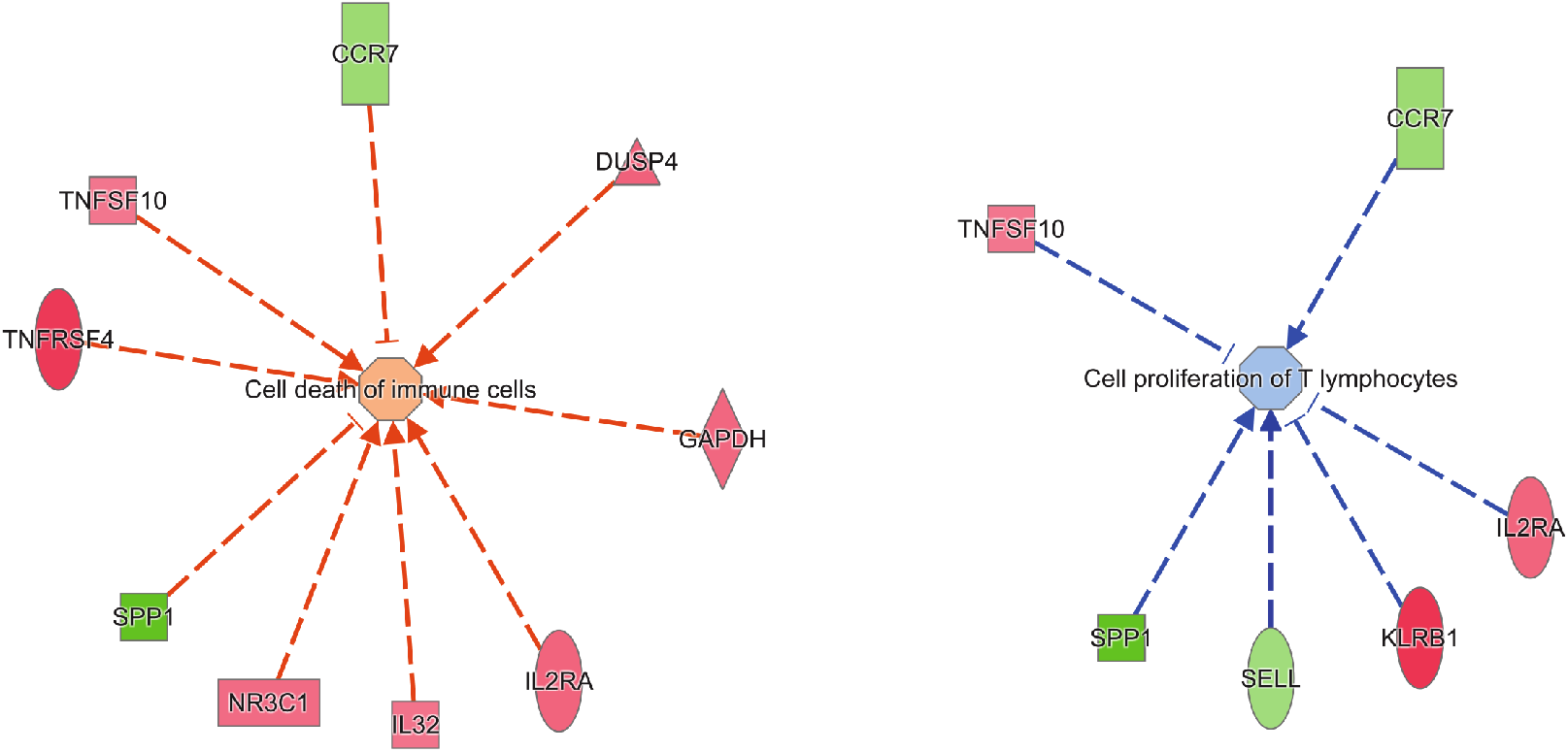
Ingenuity Pathway analysis (IPA) was used to analyze the set of genes that are differentially expressed between cells with active viral gene expression (GFP+) and those with silenced viral gene expression (GFP-). Graphical representation of two pathways enriched in this differentially expressed gene set are shown. Genes expressed at a higher level in GFP-cells are shown in green, while genes expressed at higher level in GFP+ cells are shown in red. Arrows represent positive regulation of the cellular pathway, while bars represent negative regulation.

**Supplemental Figure 2.**
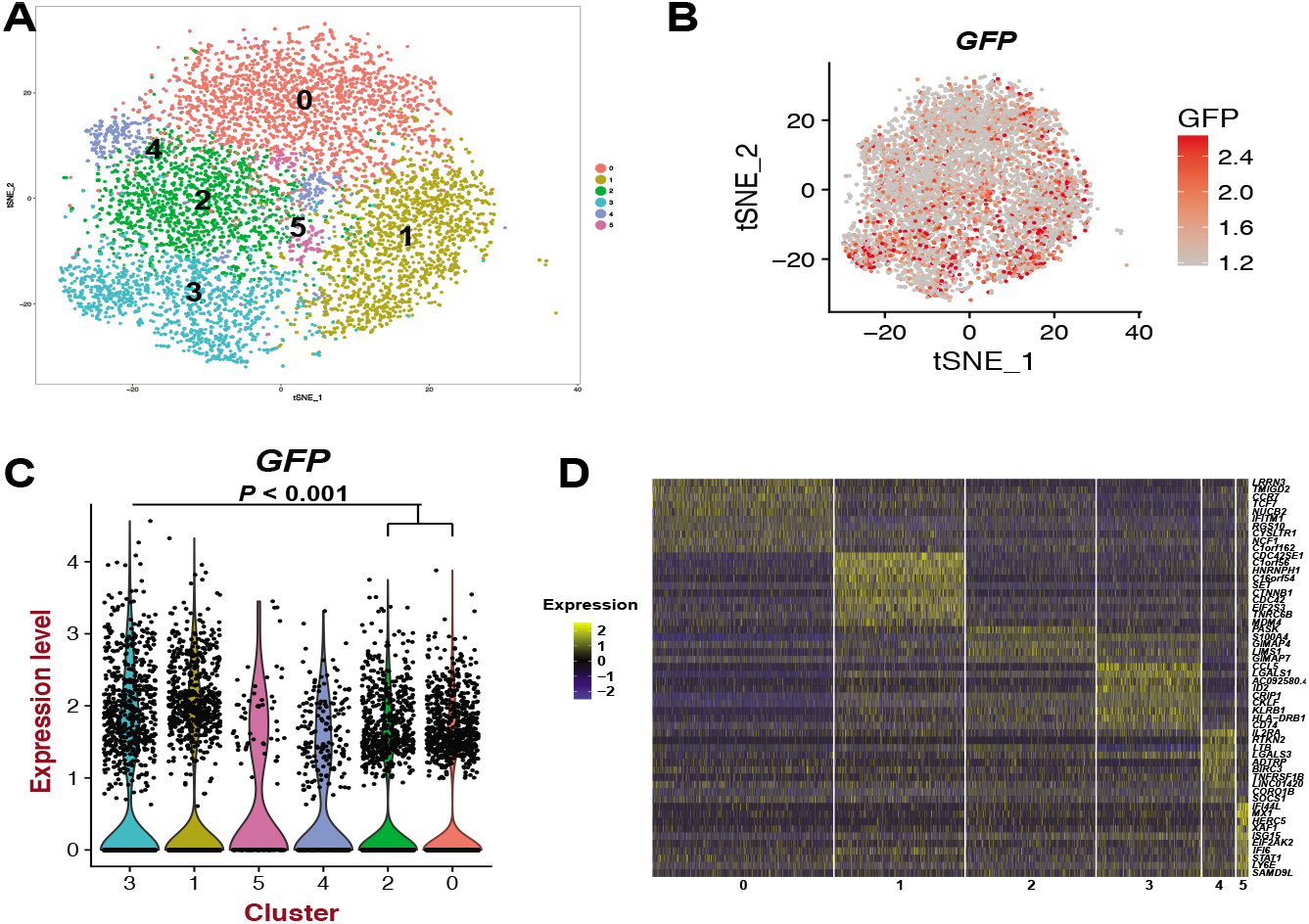
Second donor scRNAseq. **(A)**. Graph-based clustering of infected cells represented on the tSNE plot. **(B)**. Normalized transcript expression level of *GFP* on the tSNE plot representation. Legends indicate normalized transcript values. **(C)**. Violin plots of *GFP* expression within the identified clusters. **(D)**. Heatmap of the top 10 transcripts that define each cluster. Panels represent individual clusters and each pixel column is the expression of individual cells. Transcripts are on the rows with normalized expression (Z-scores) colored by the legend (yellow, more upregulated; blue, more downregulated).

**Supplemental Table 1.**
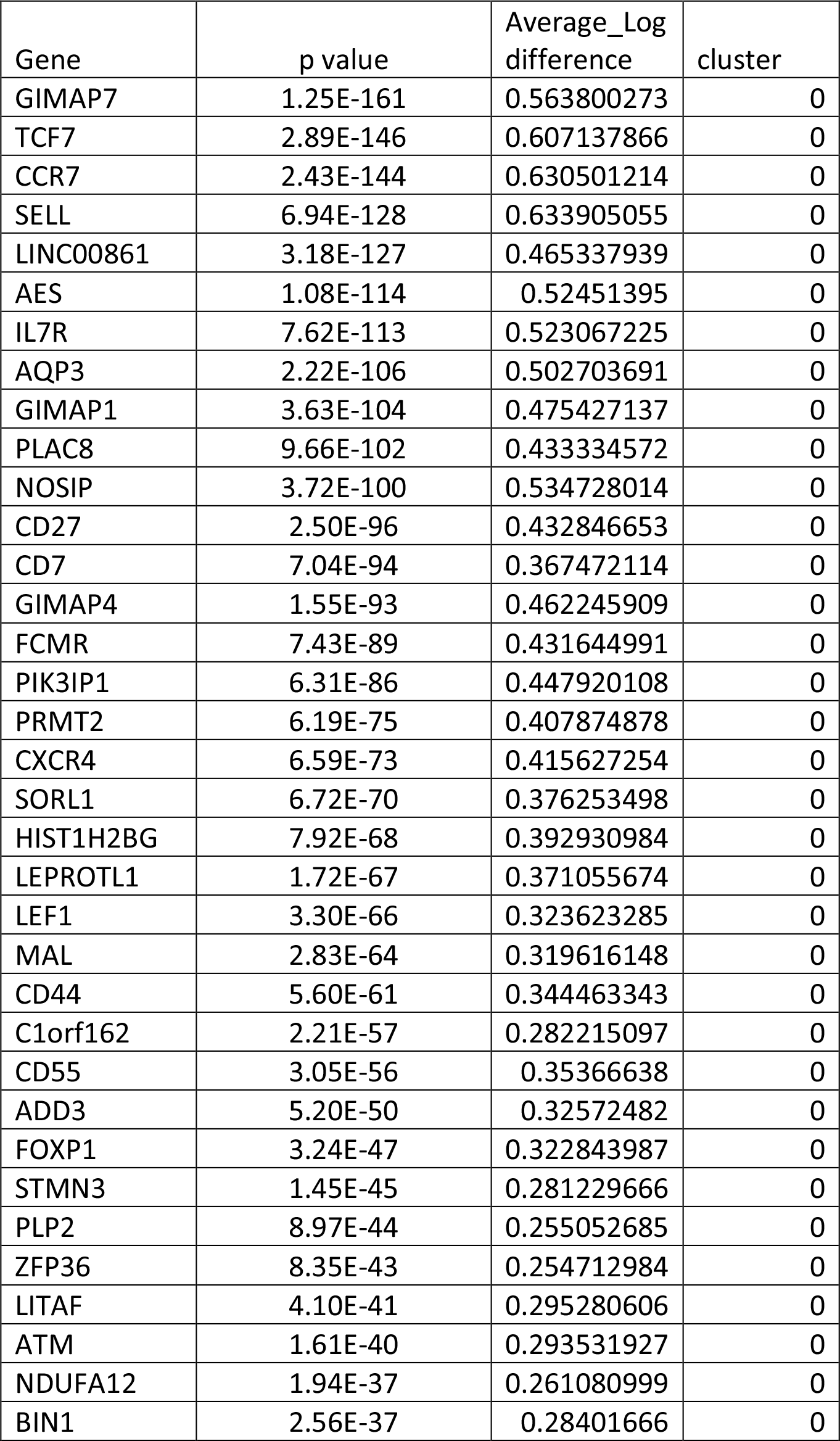
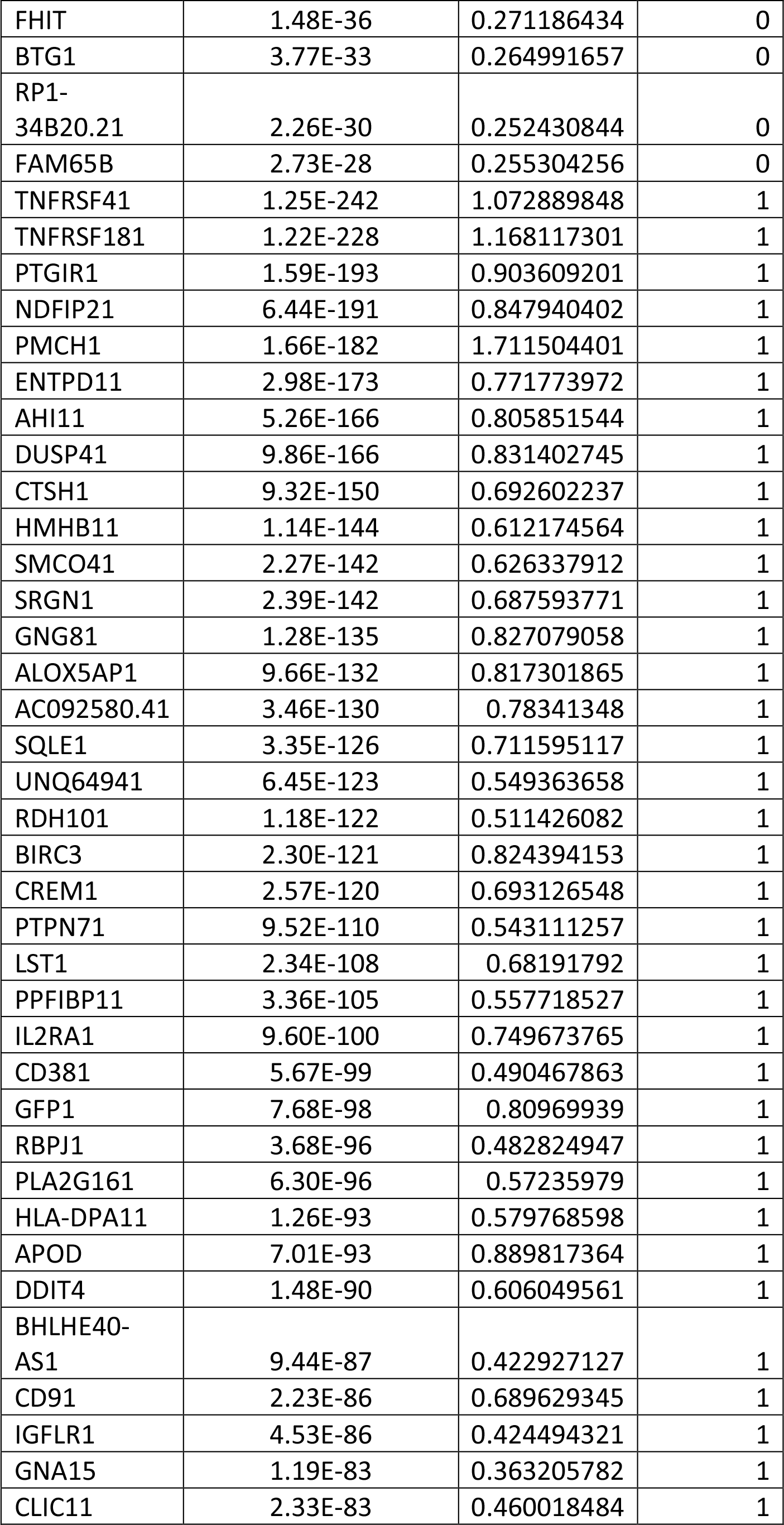
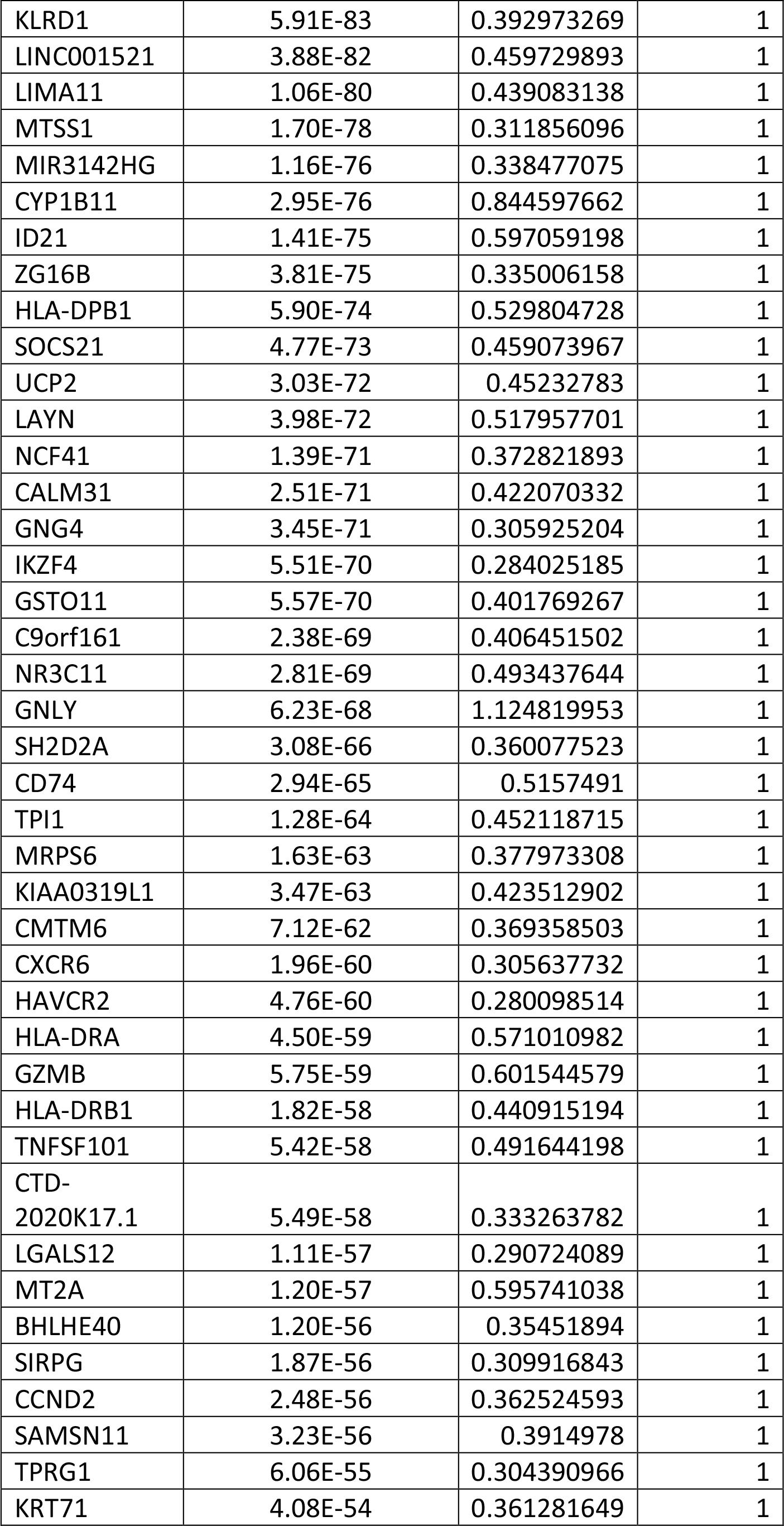
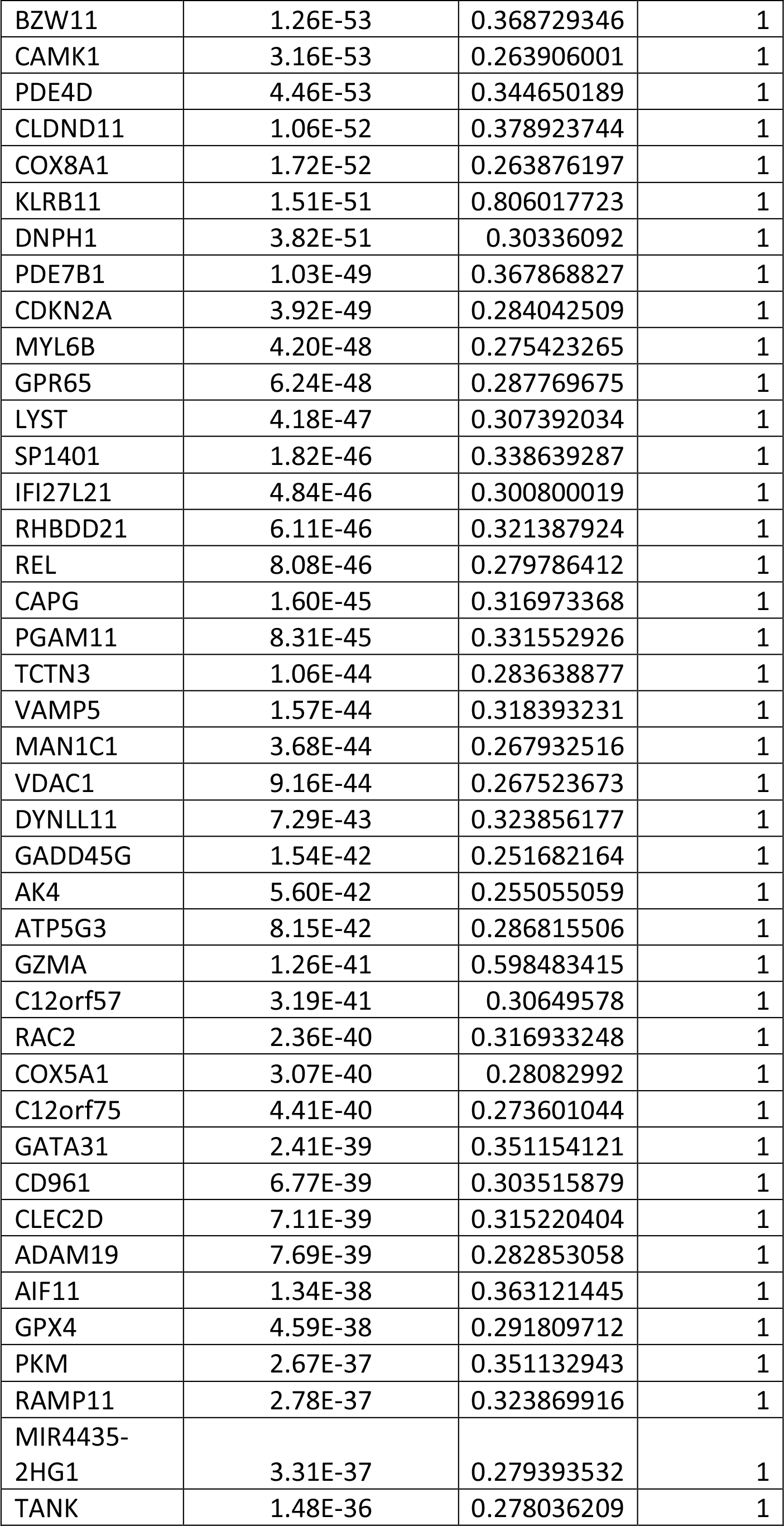
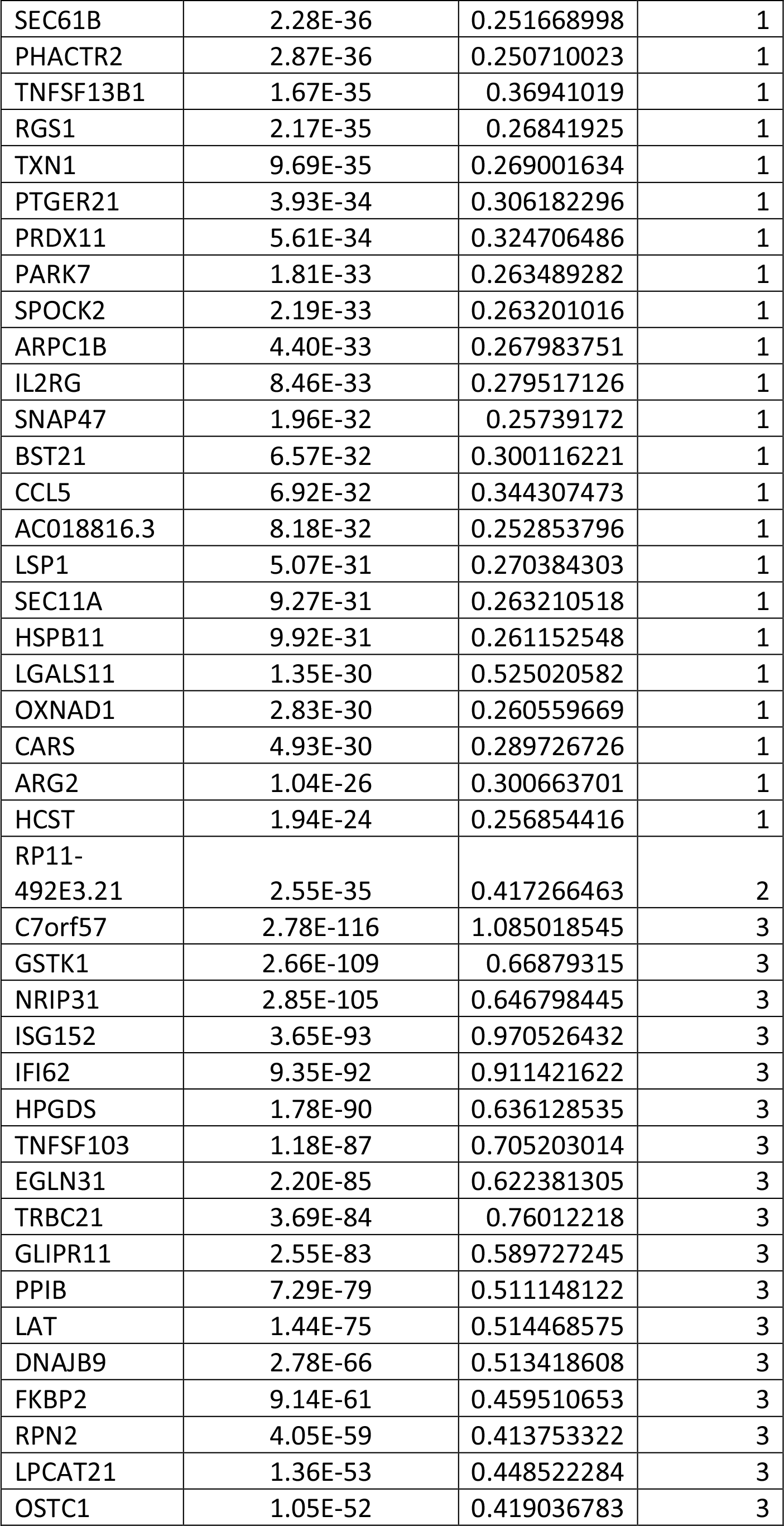
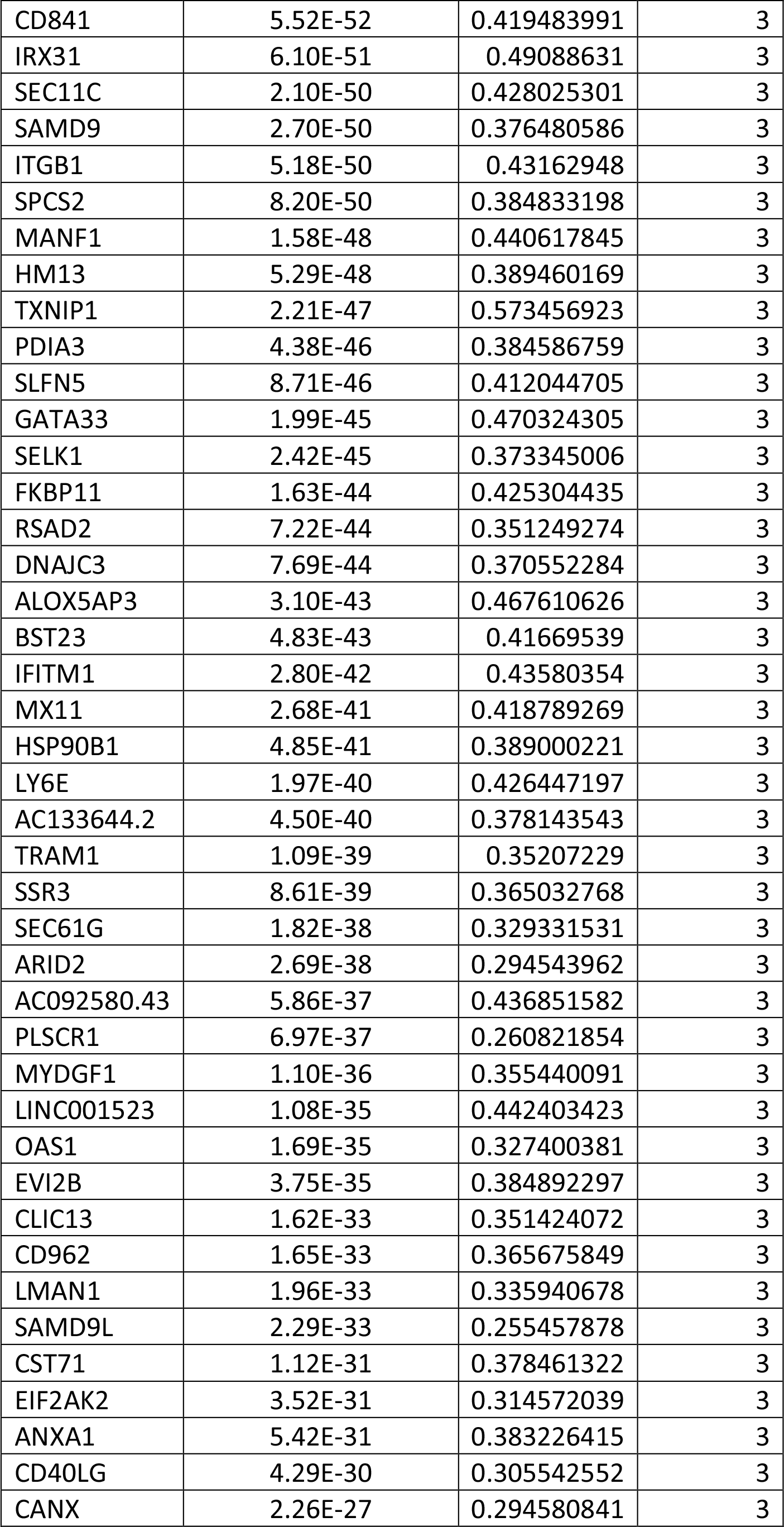
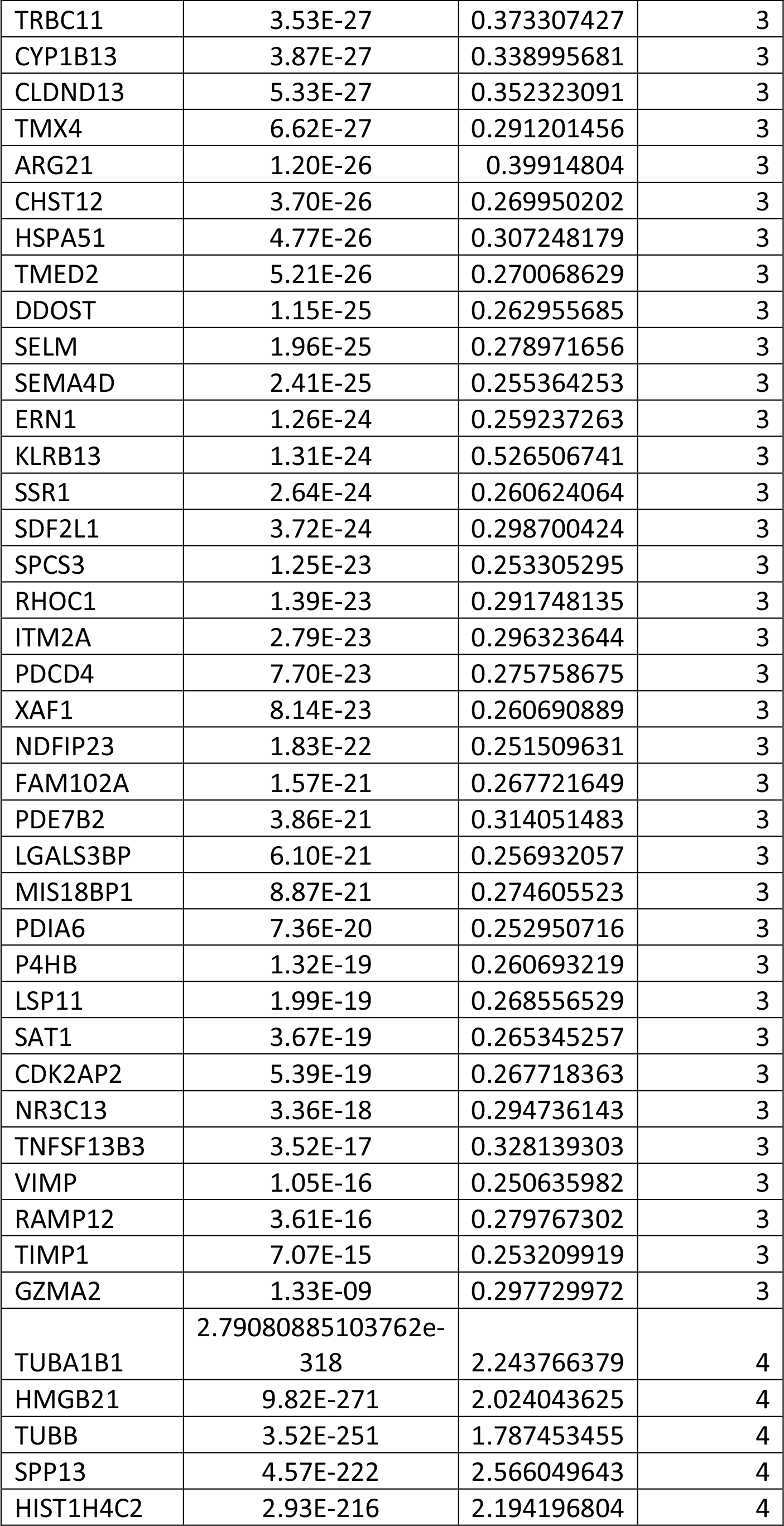
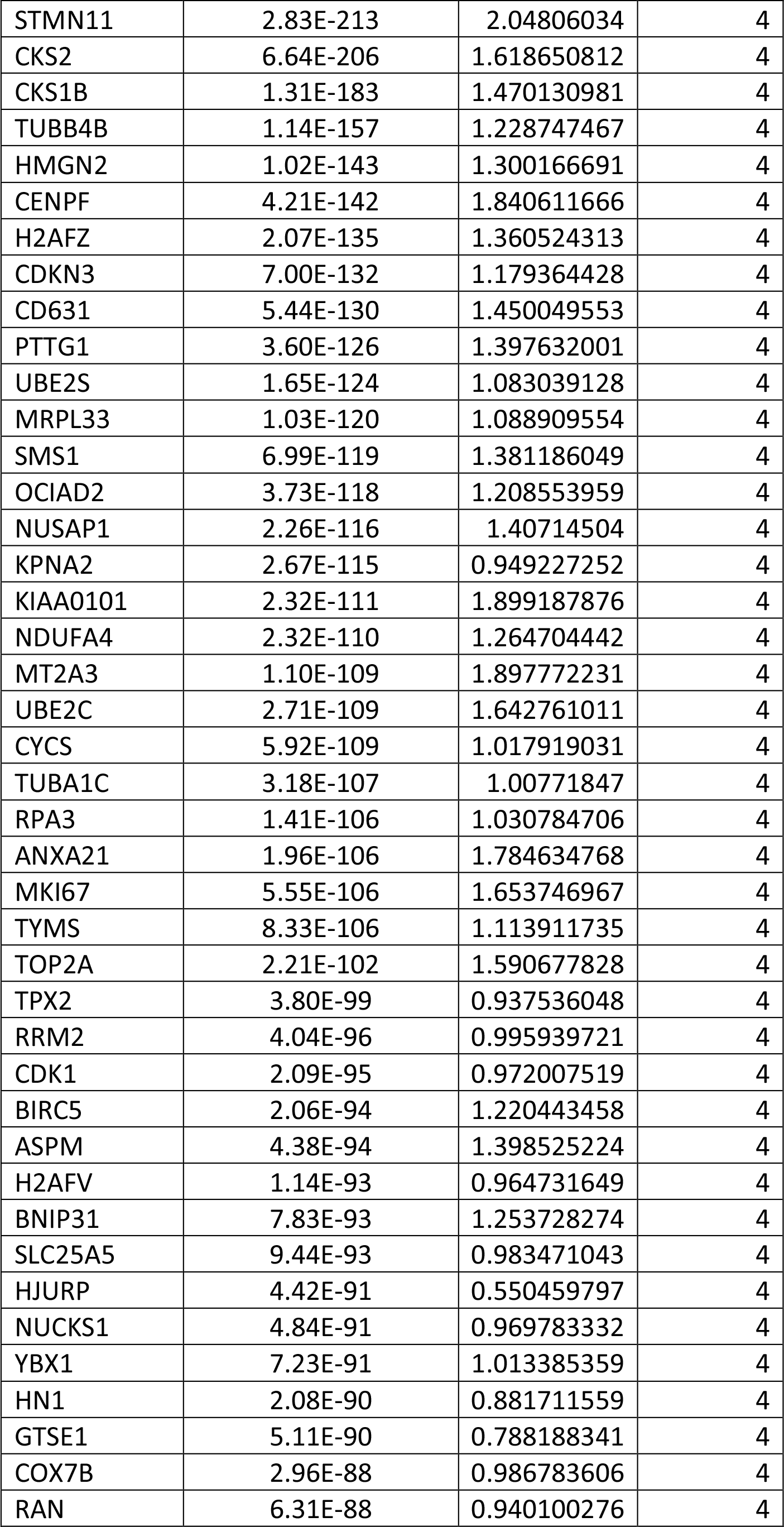
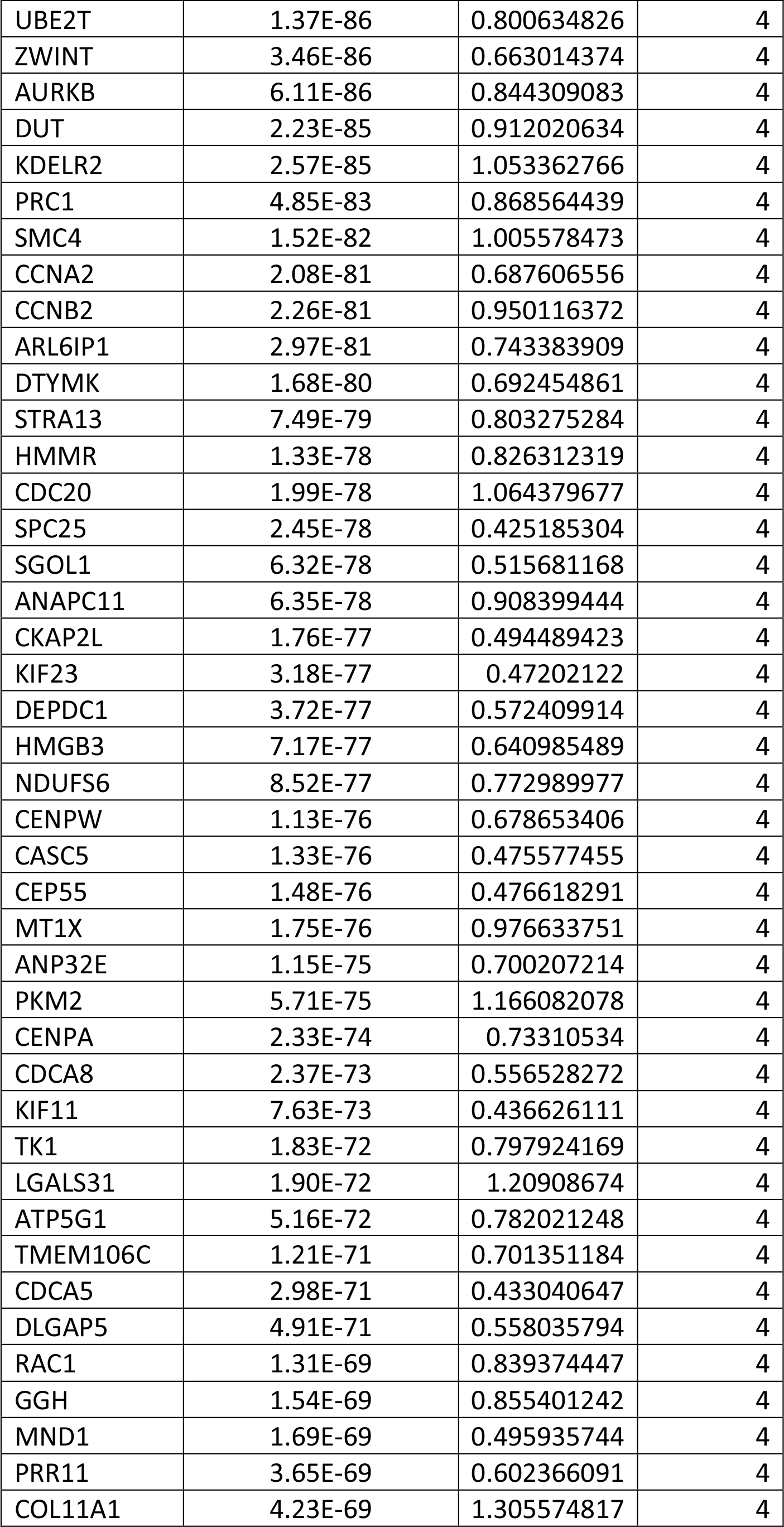
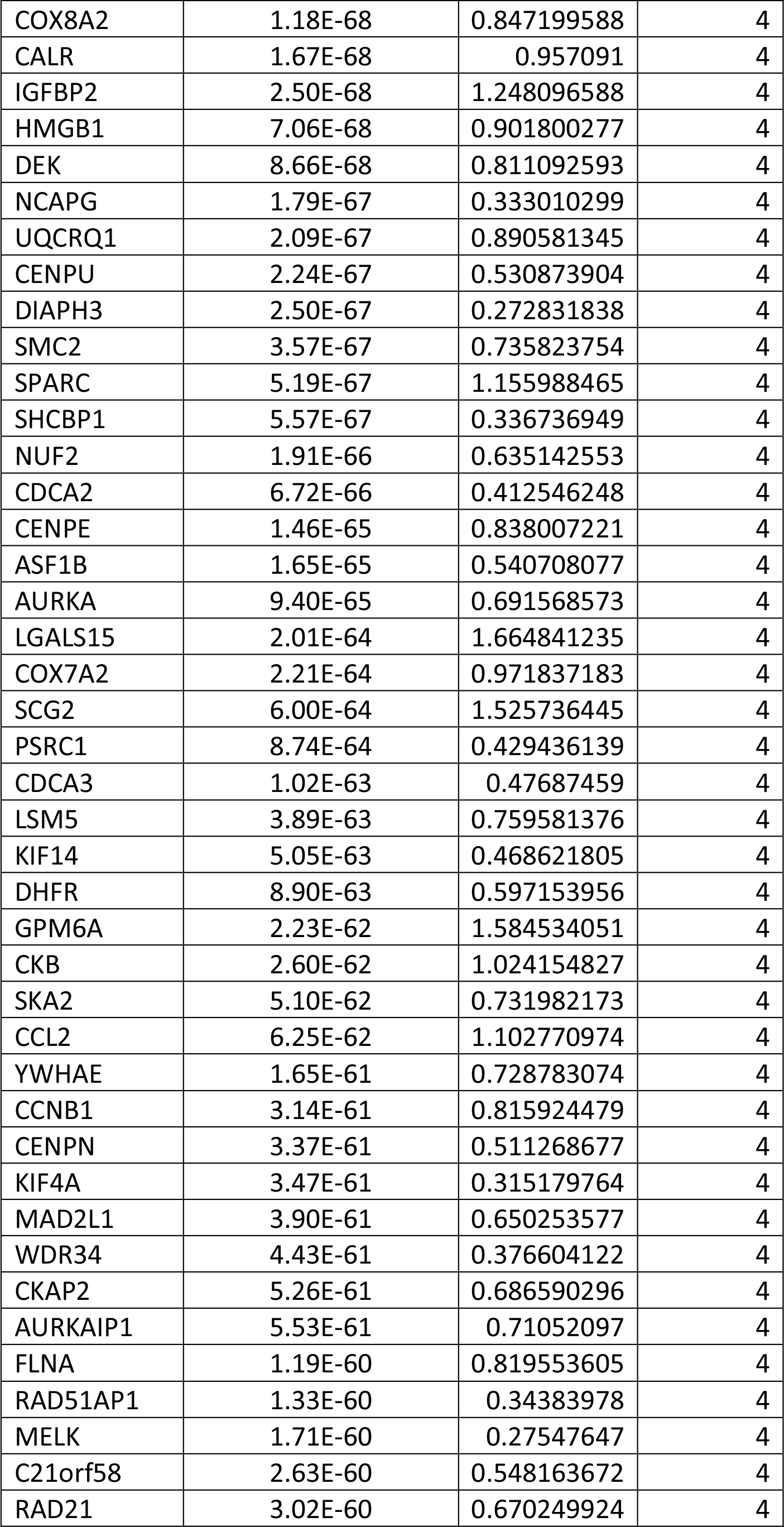
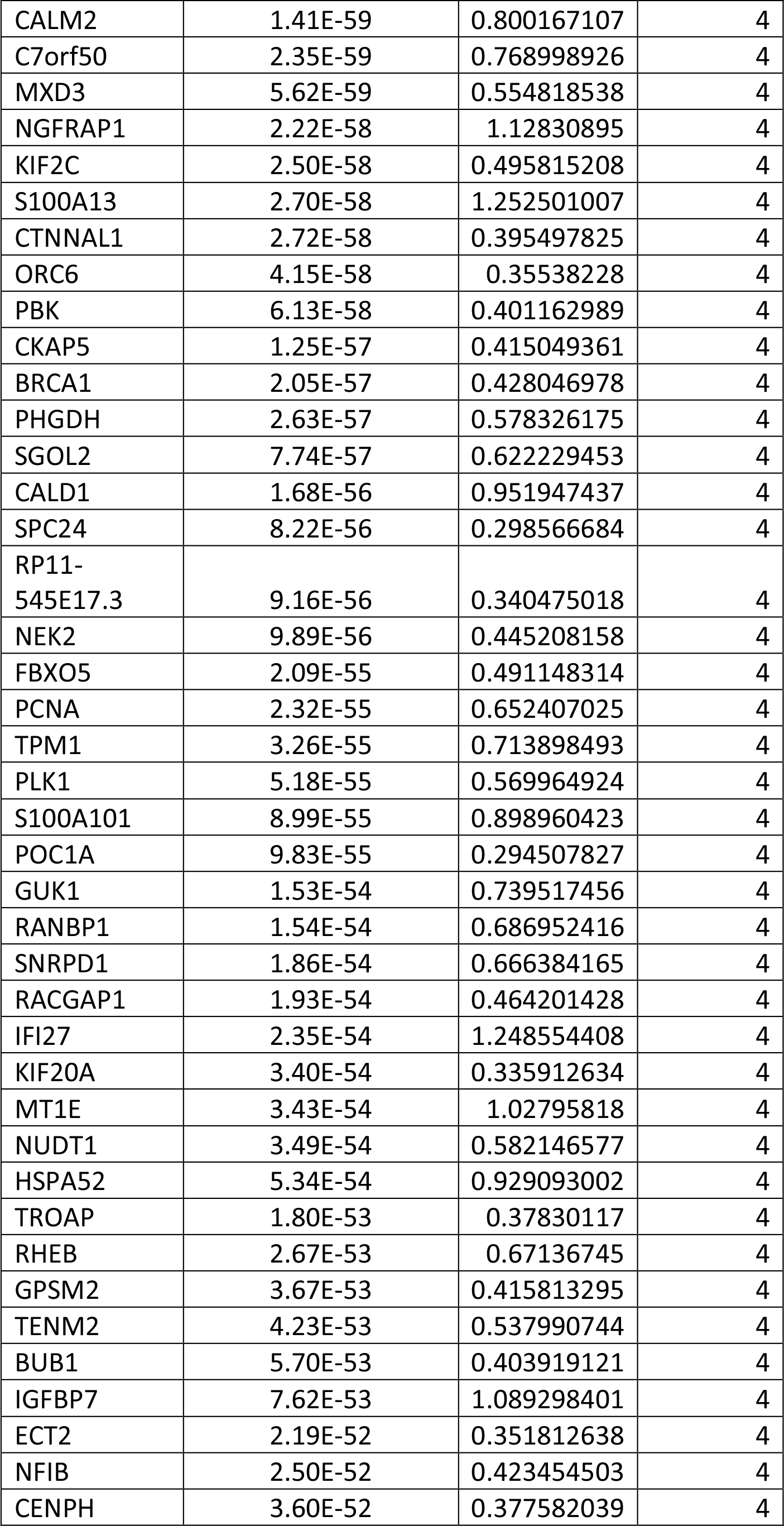
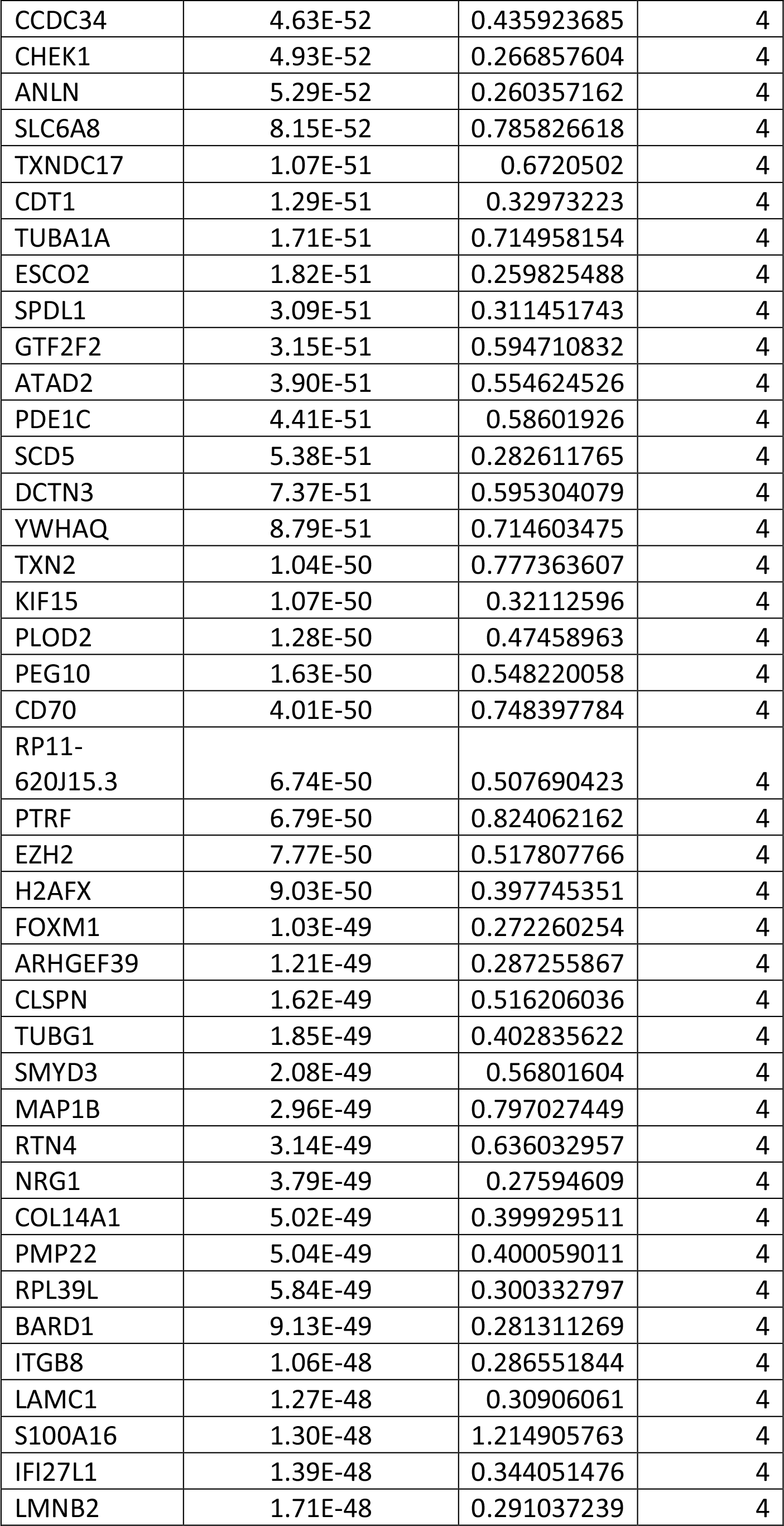
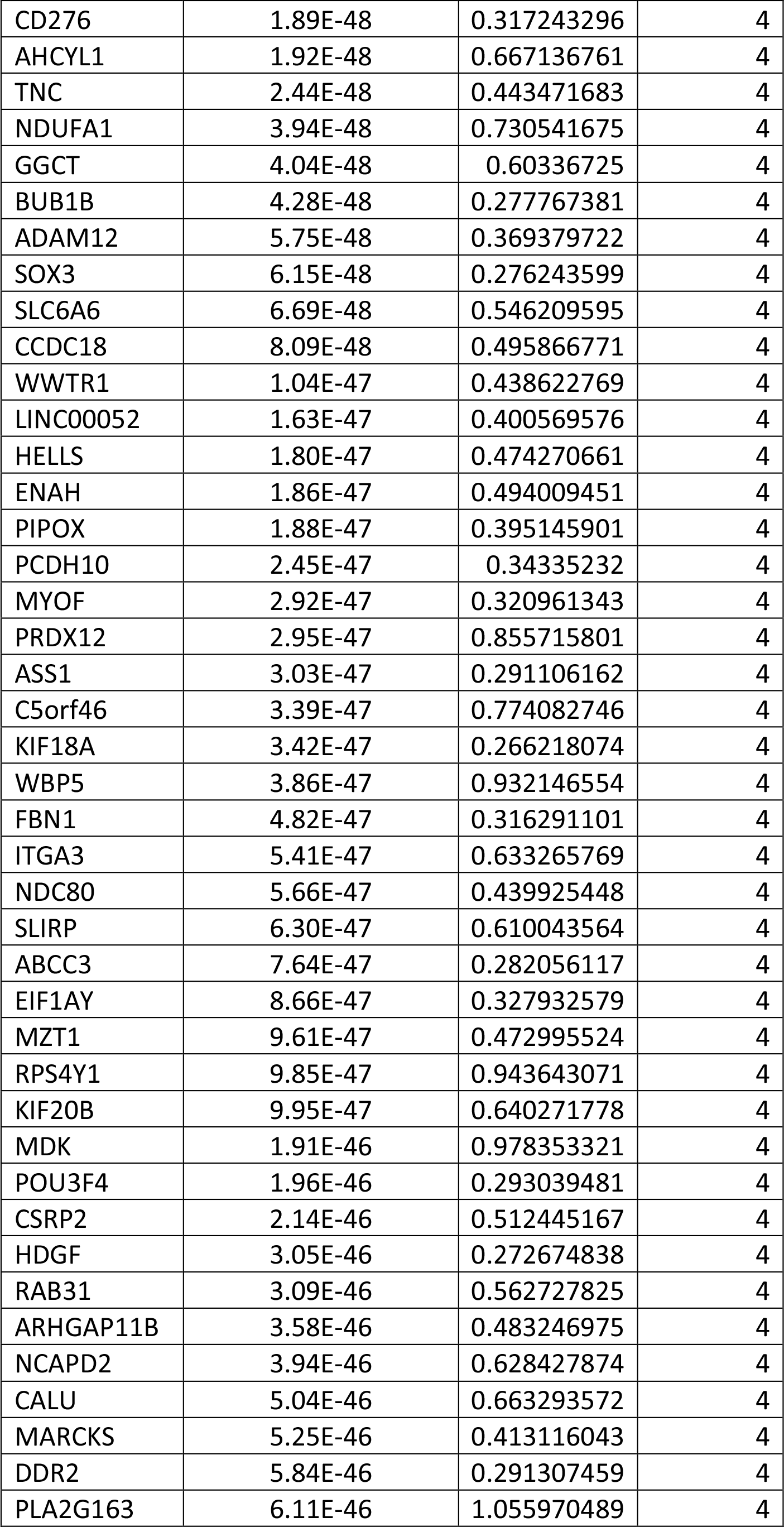
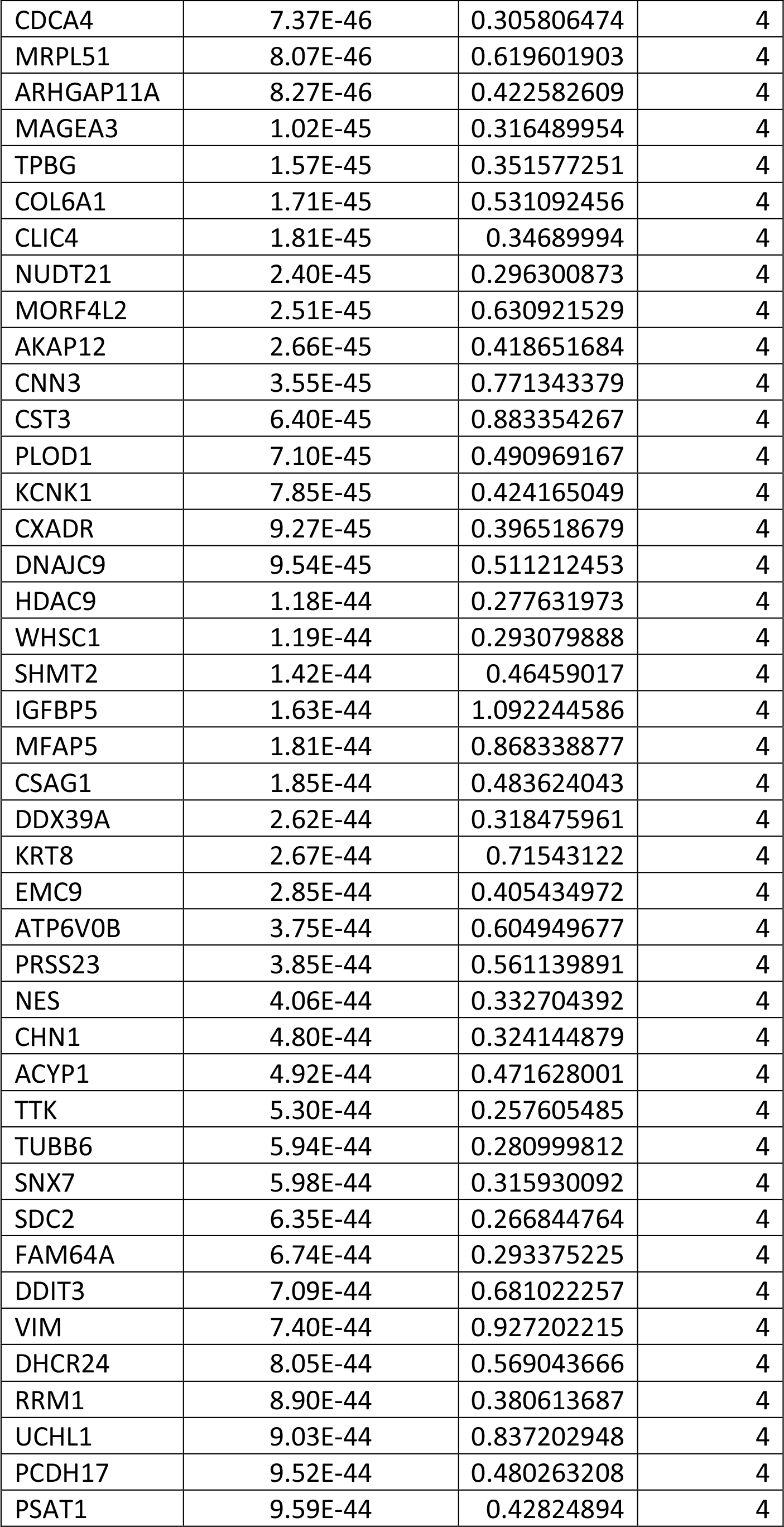
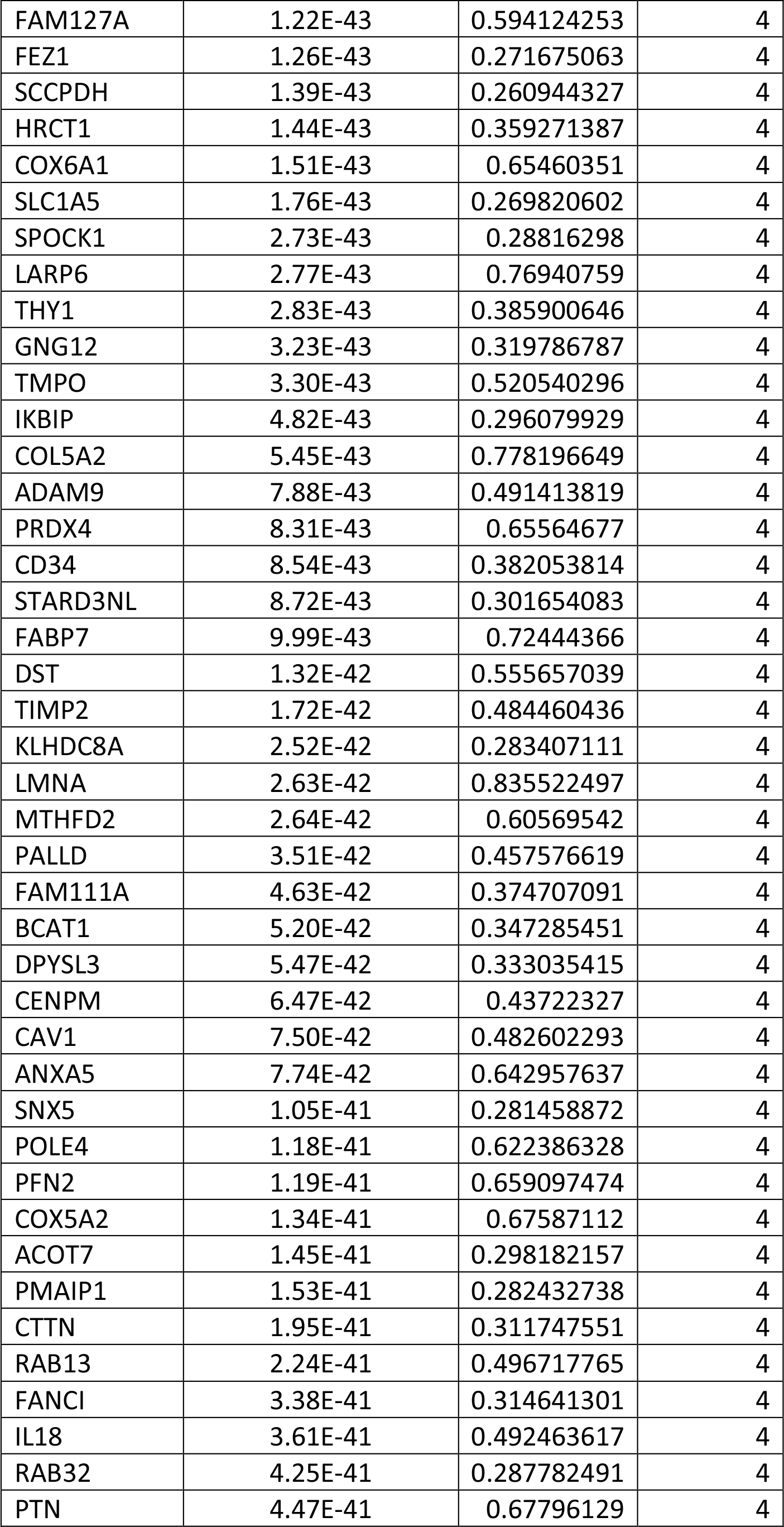
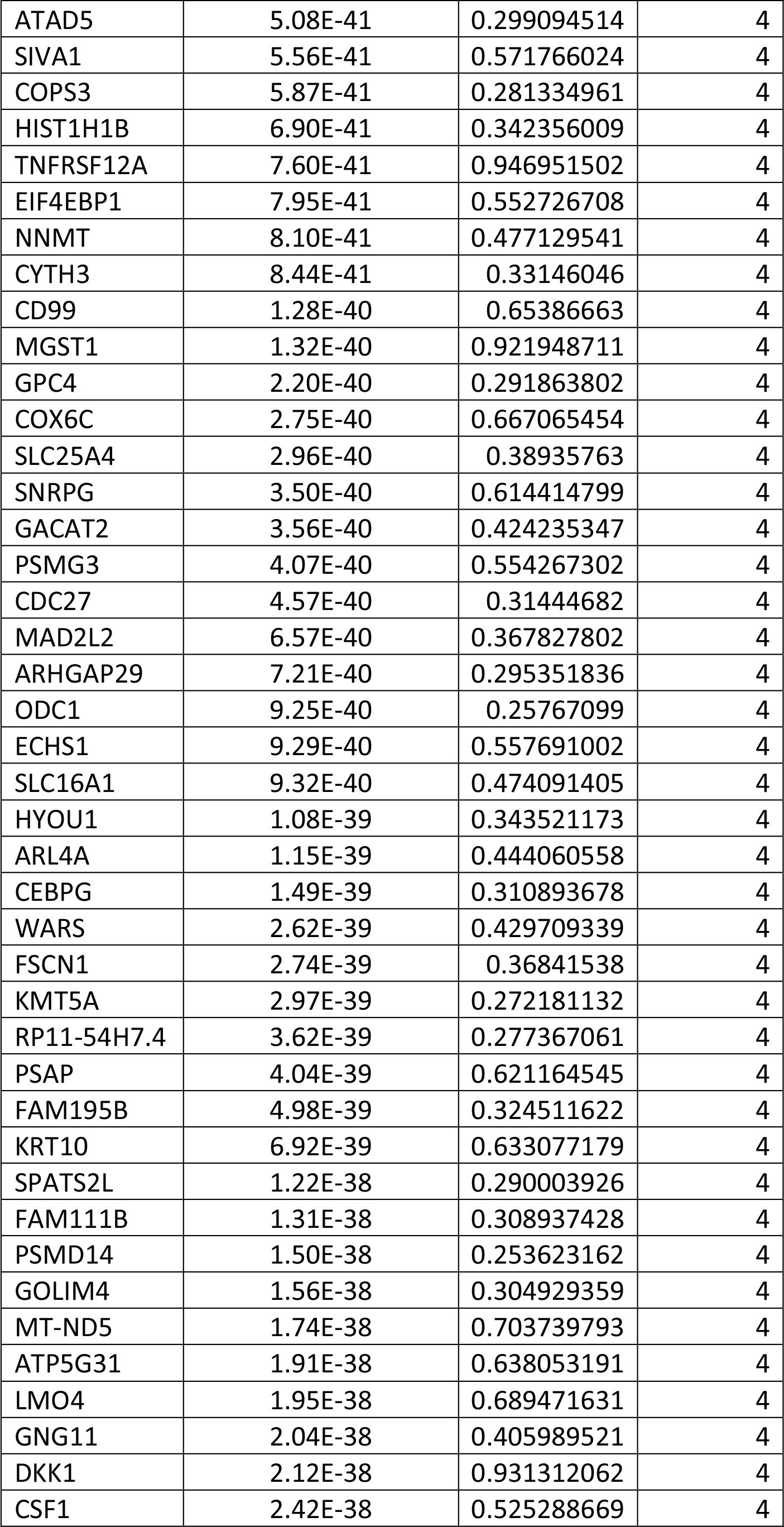
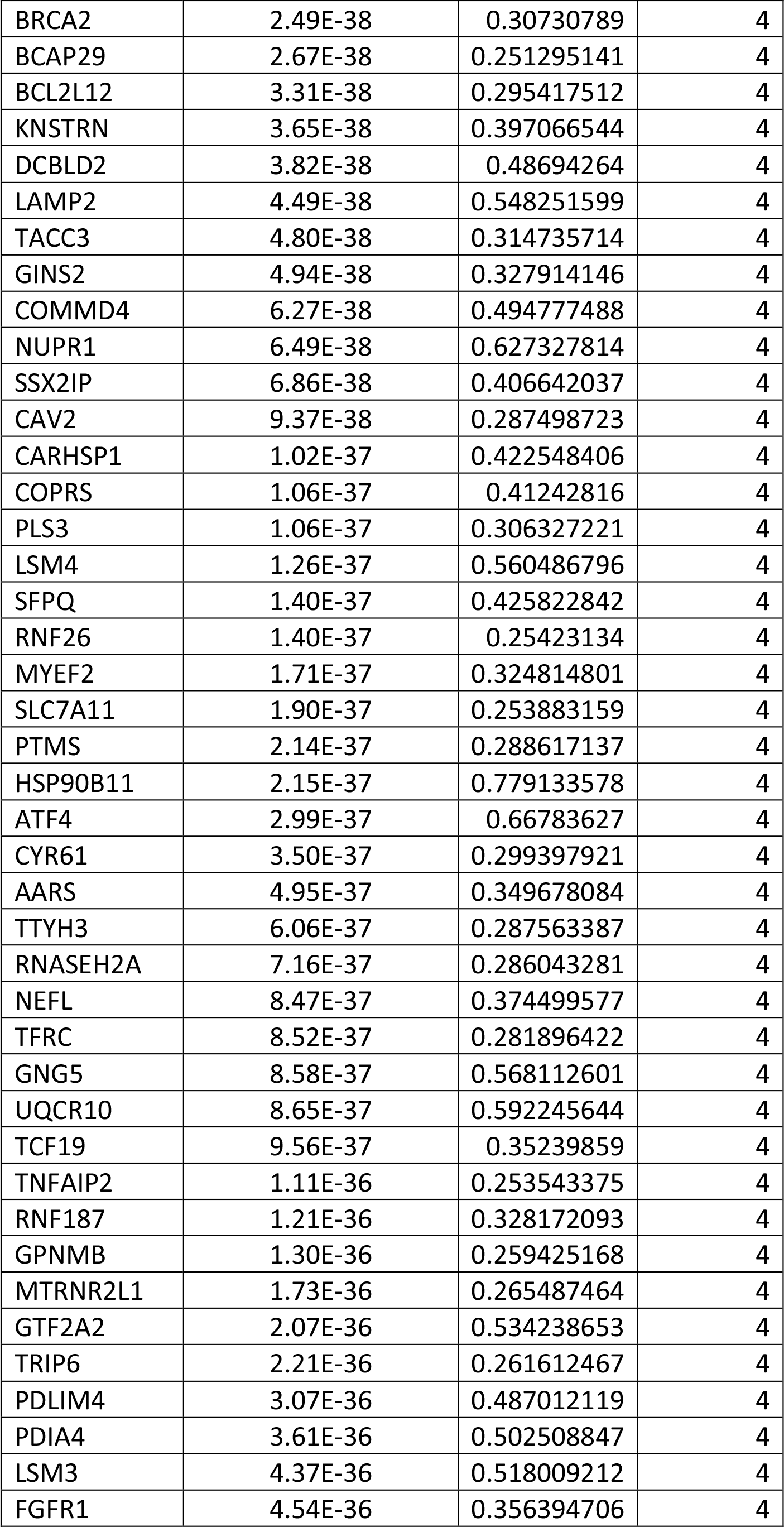
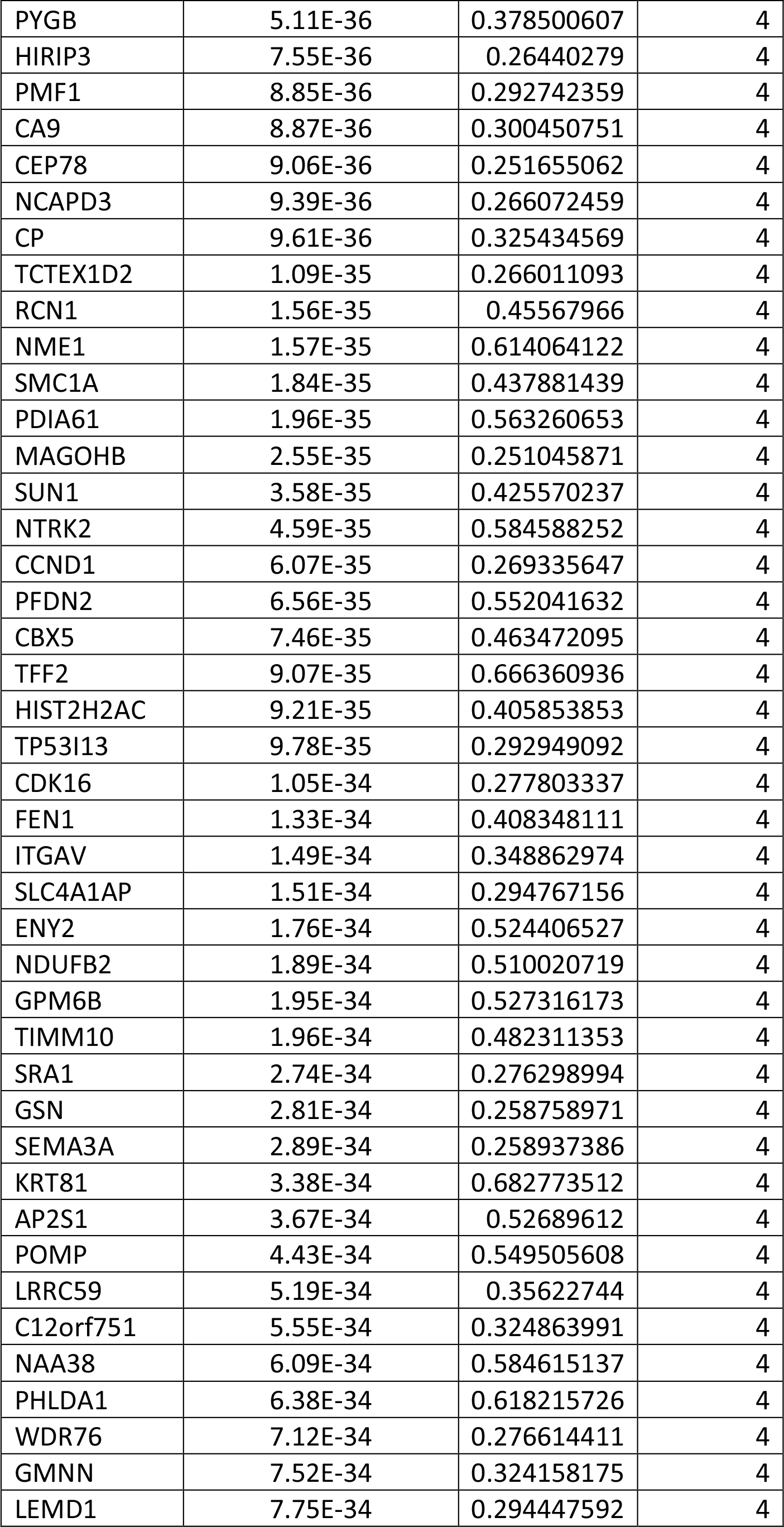
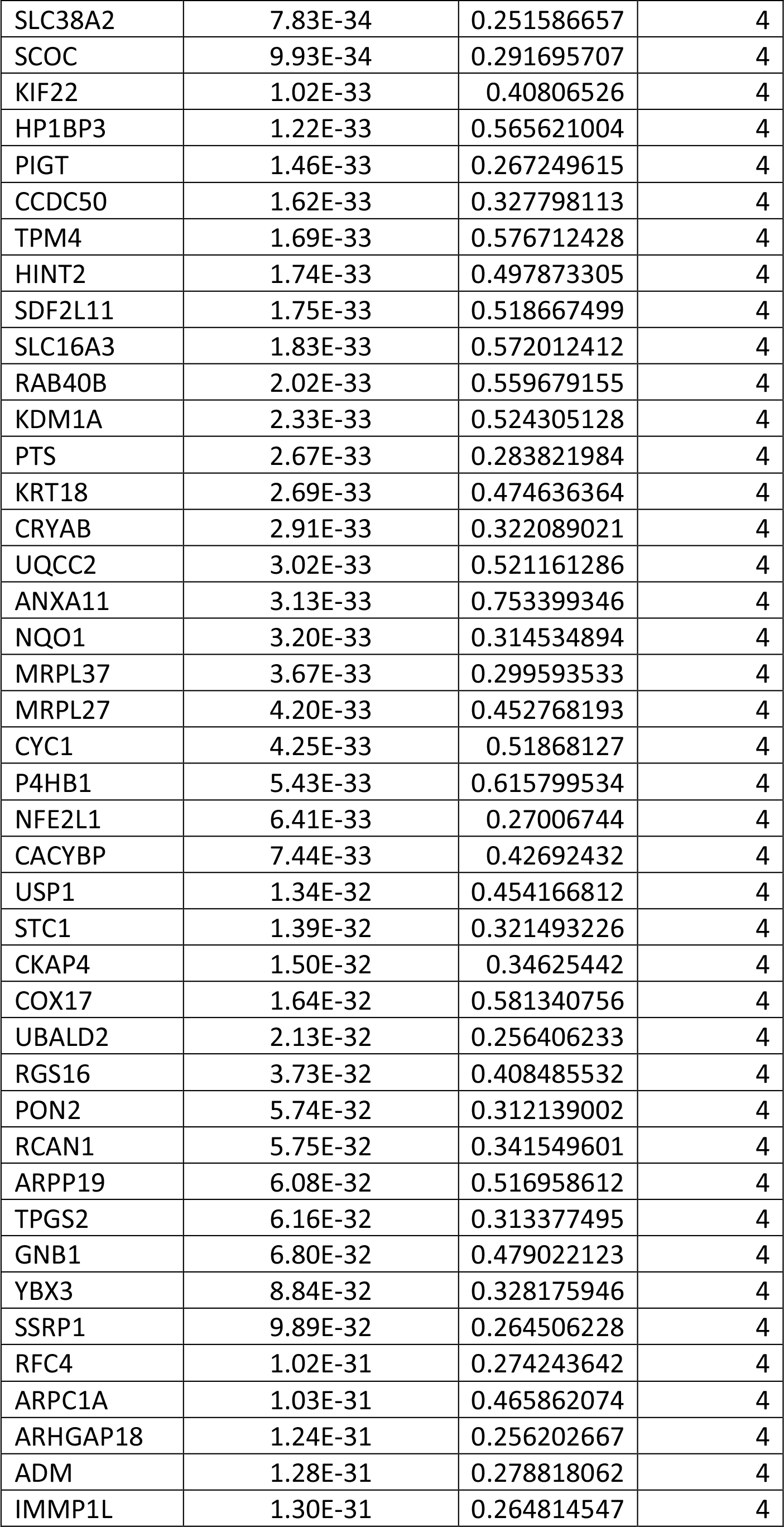
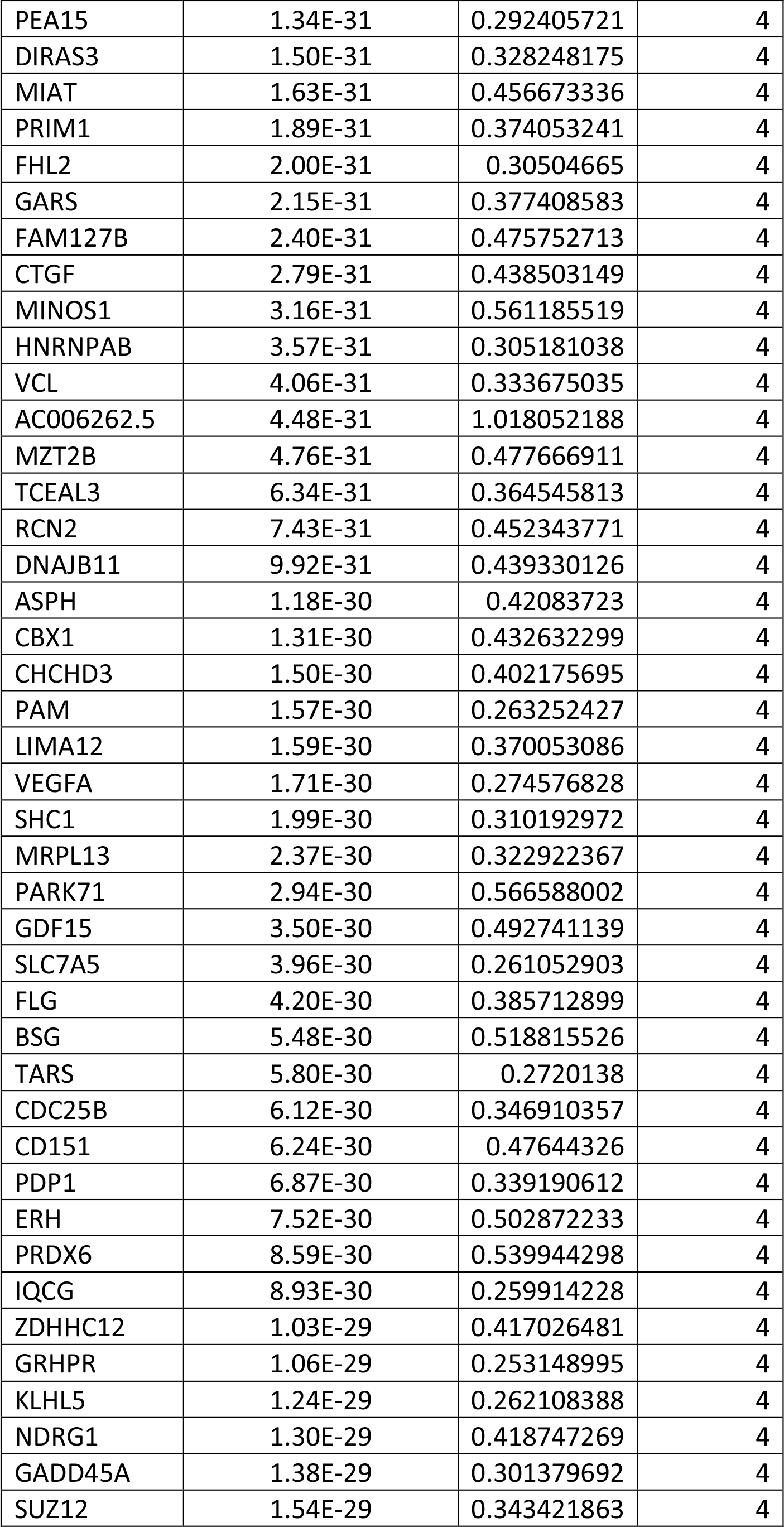
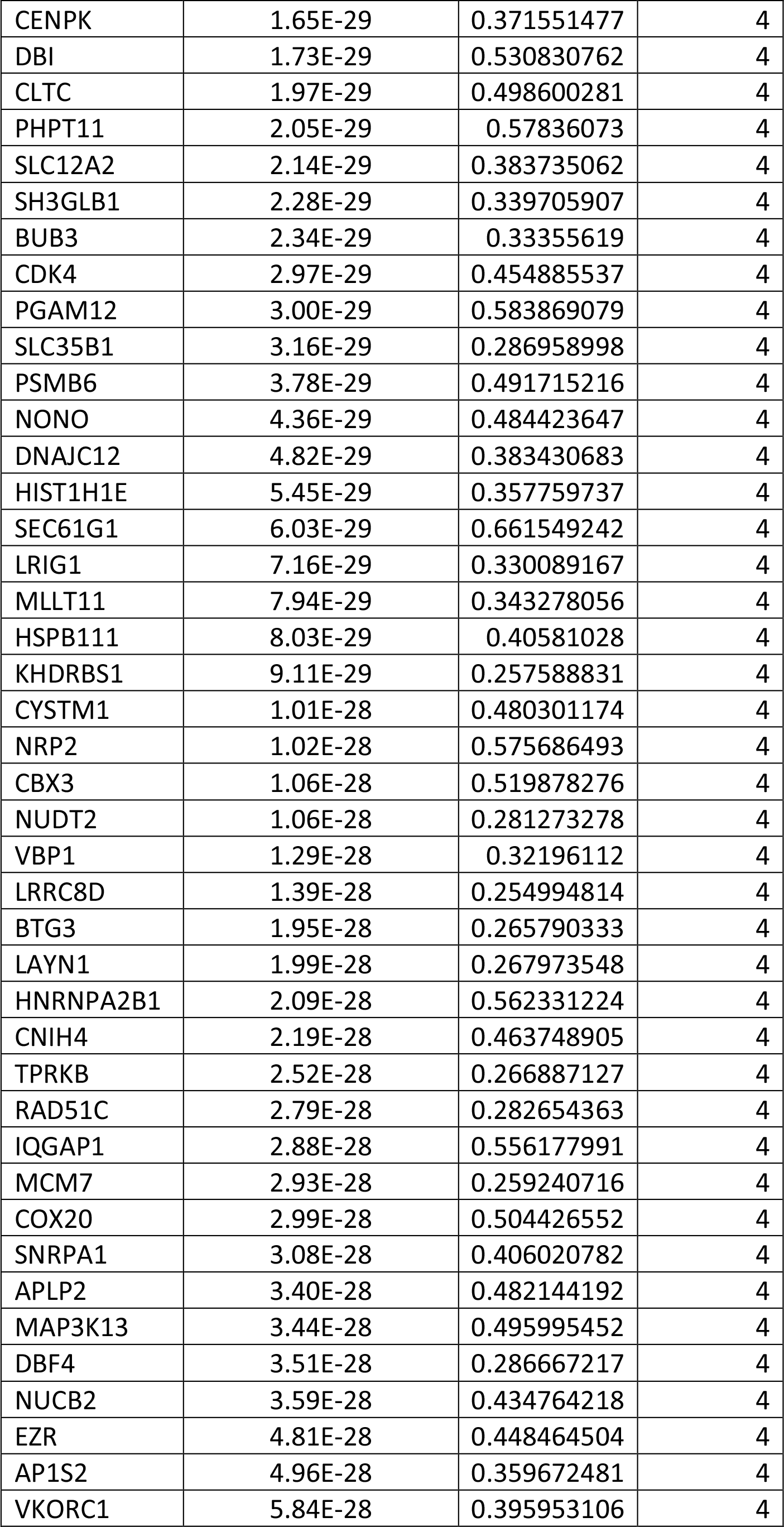
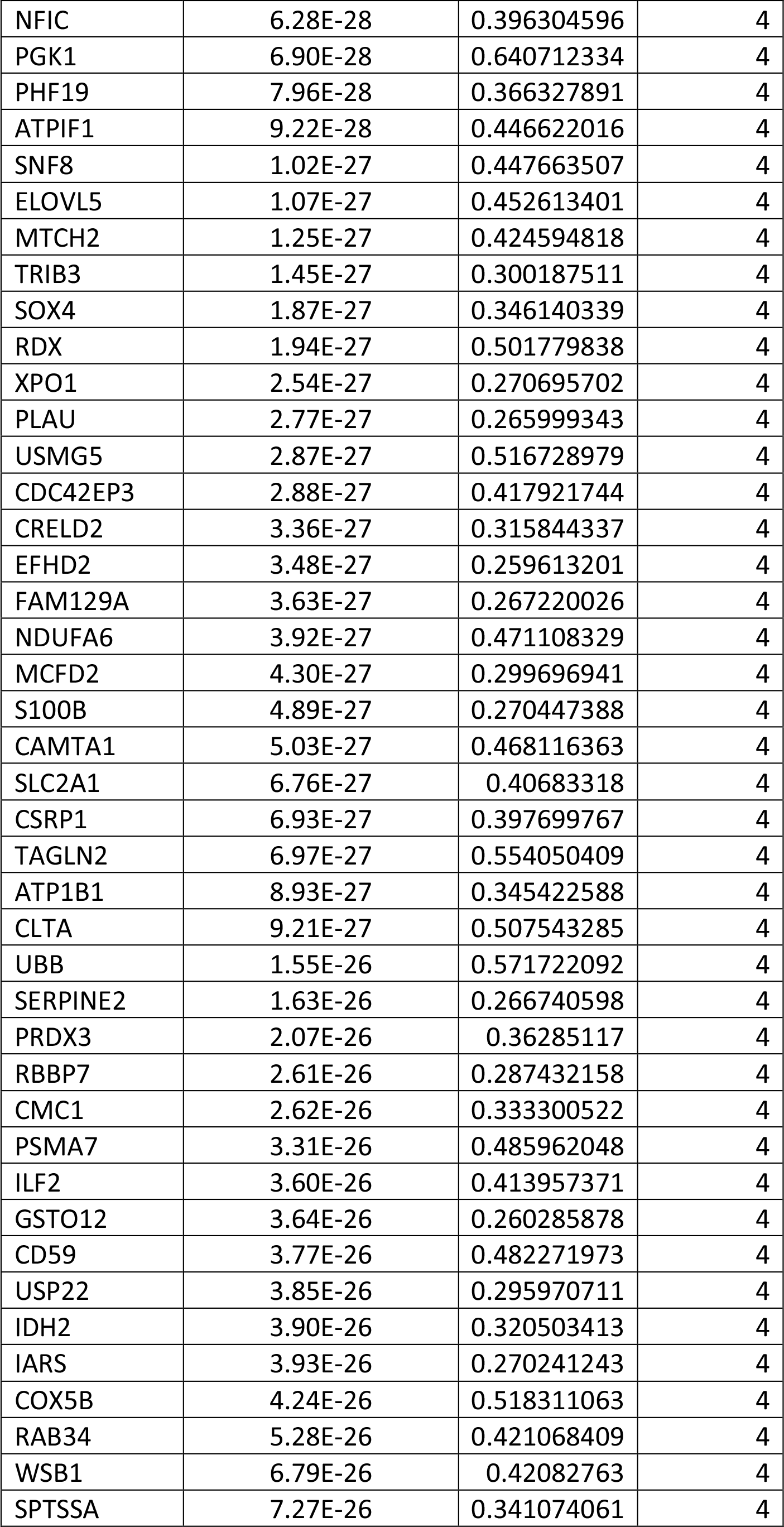
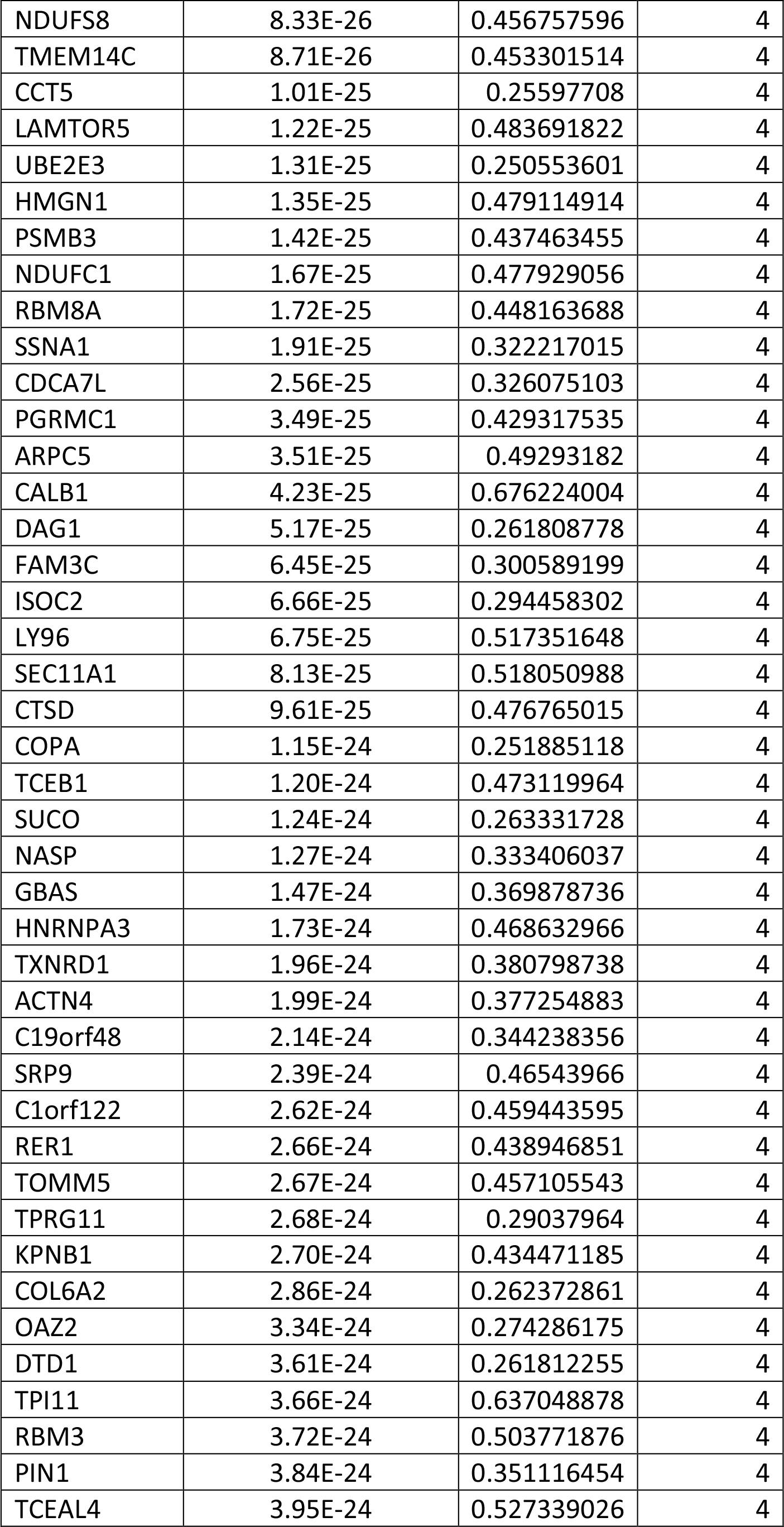
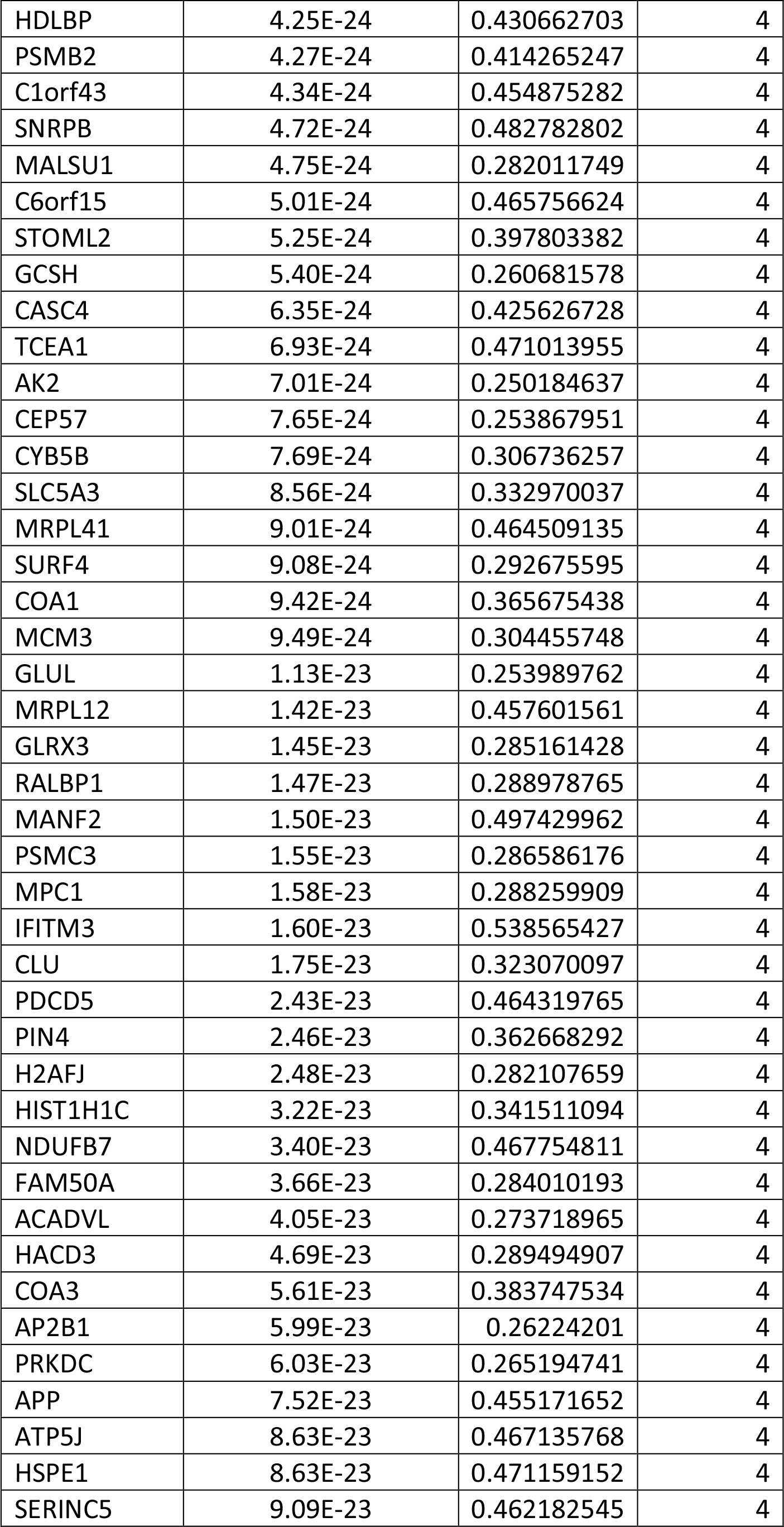
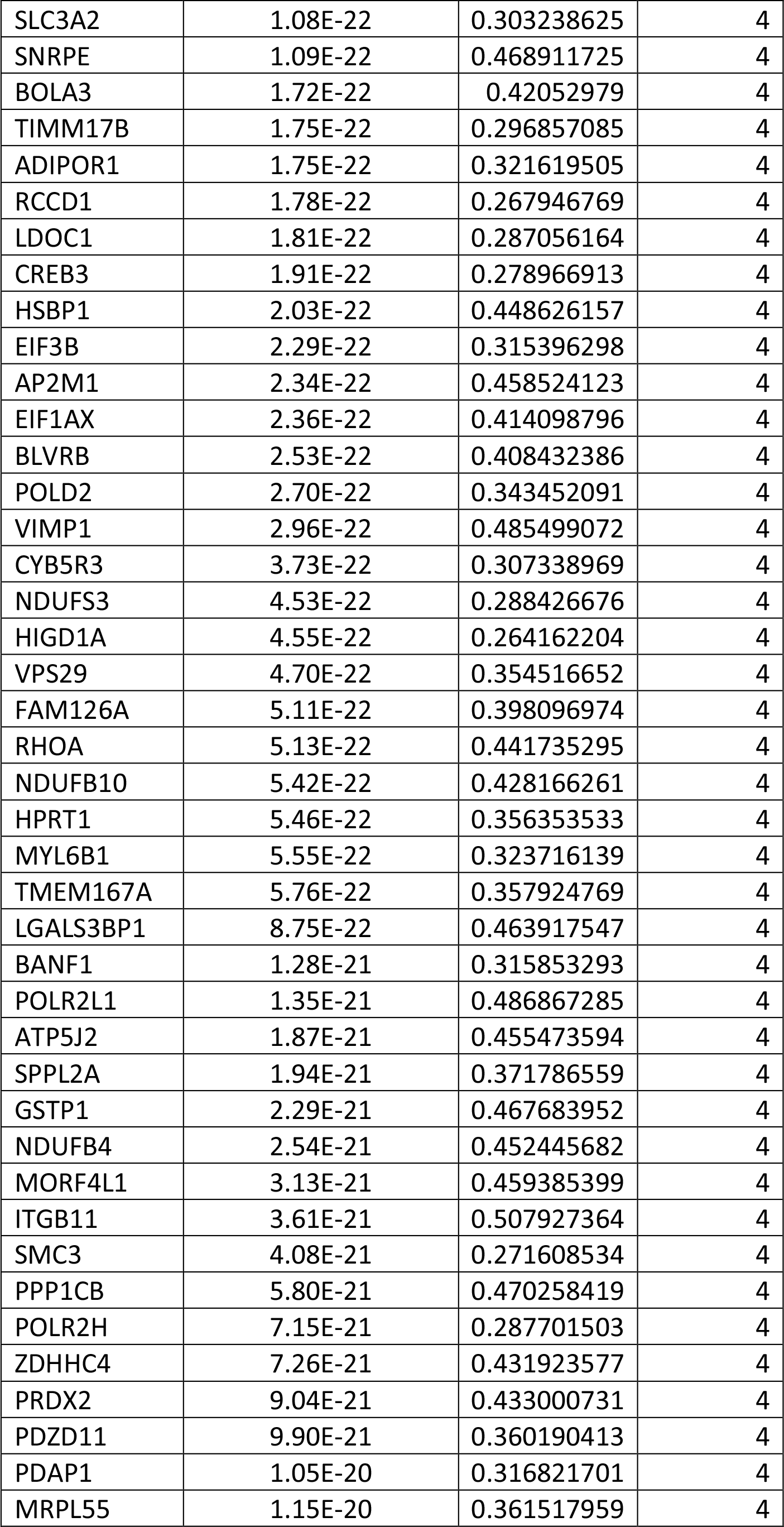
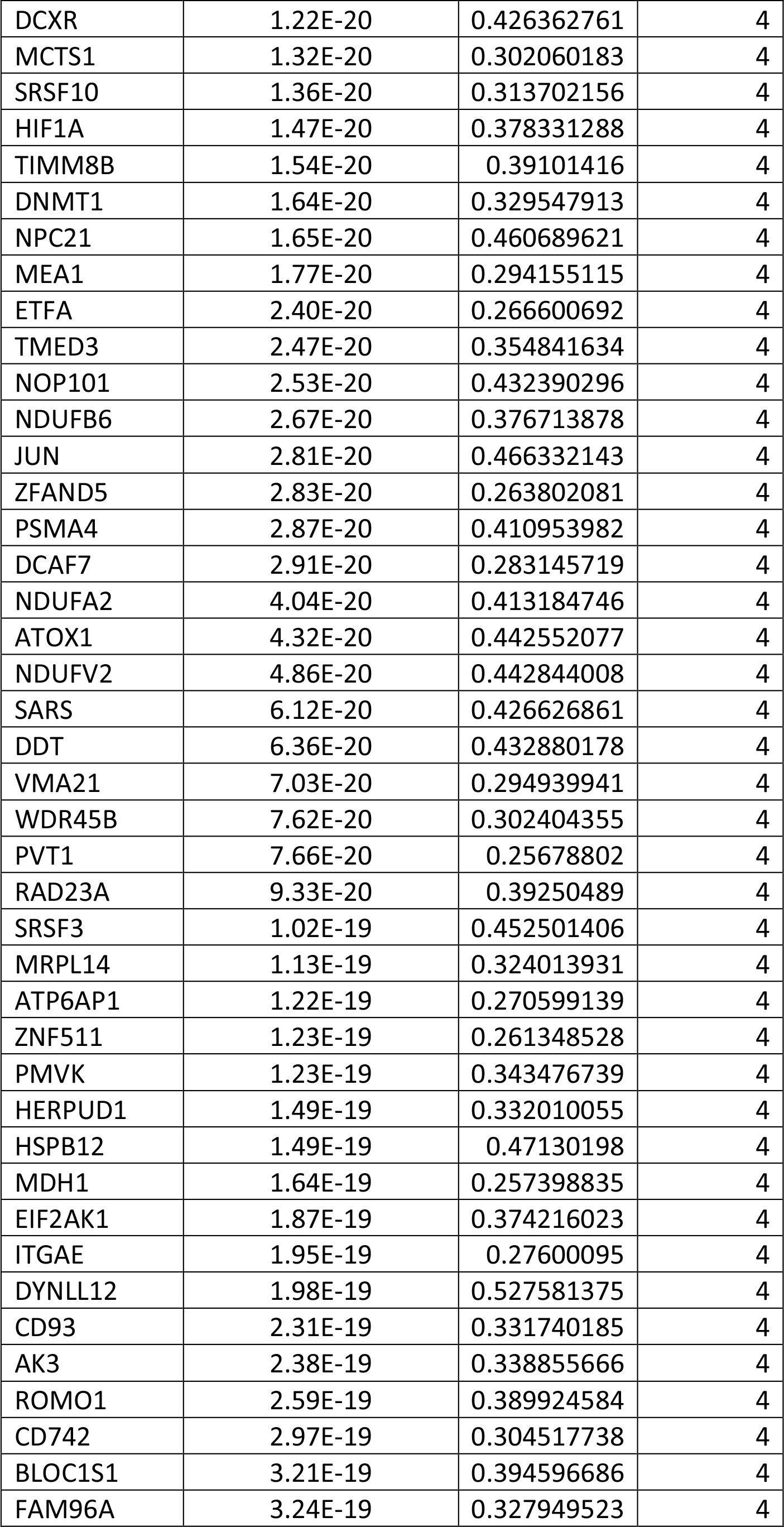
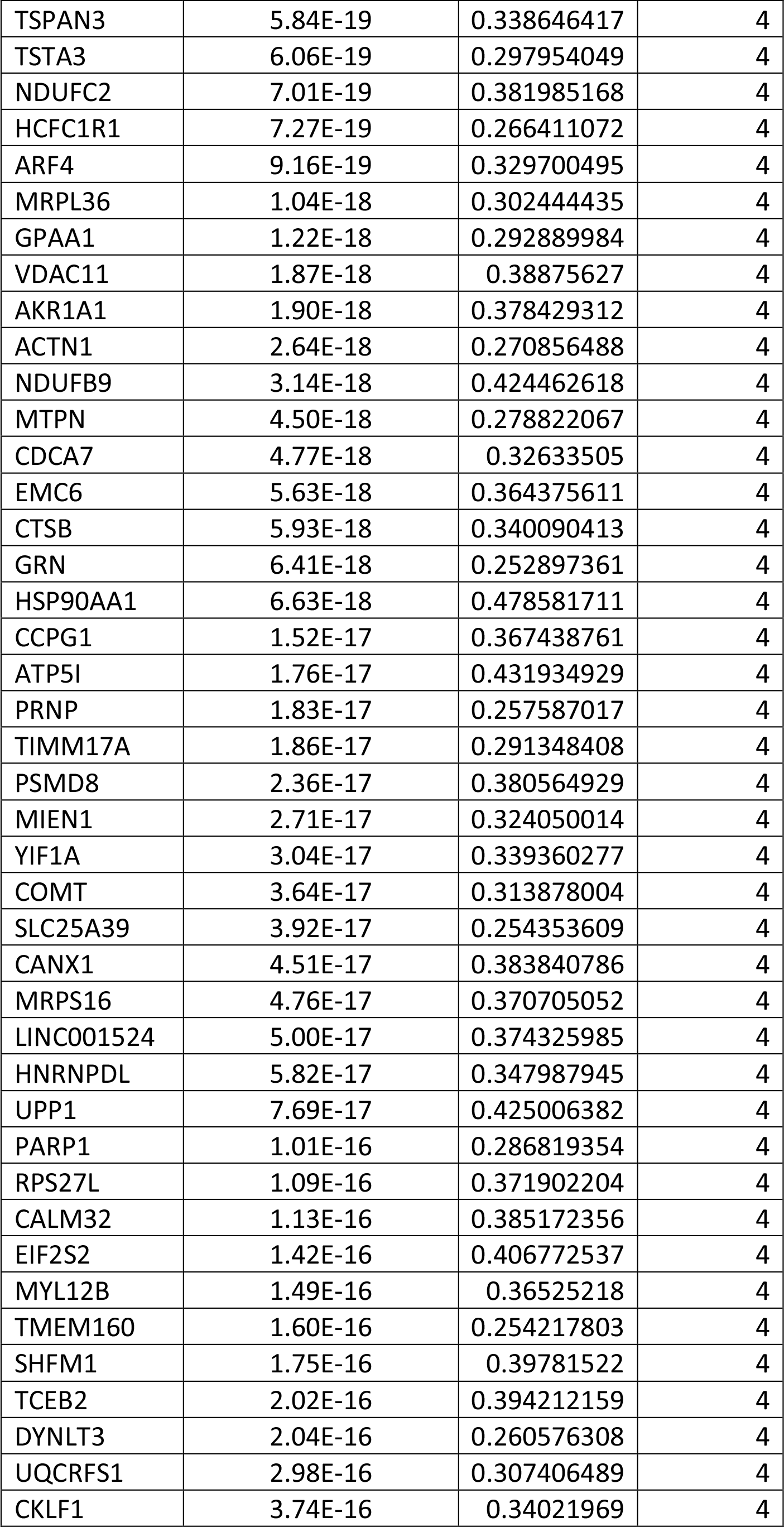
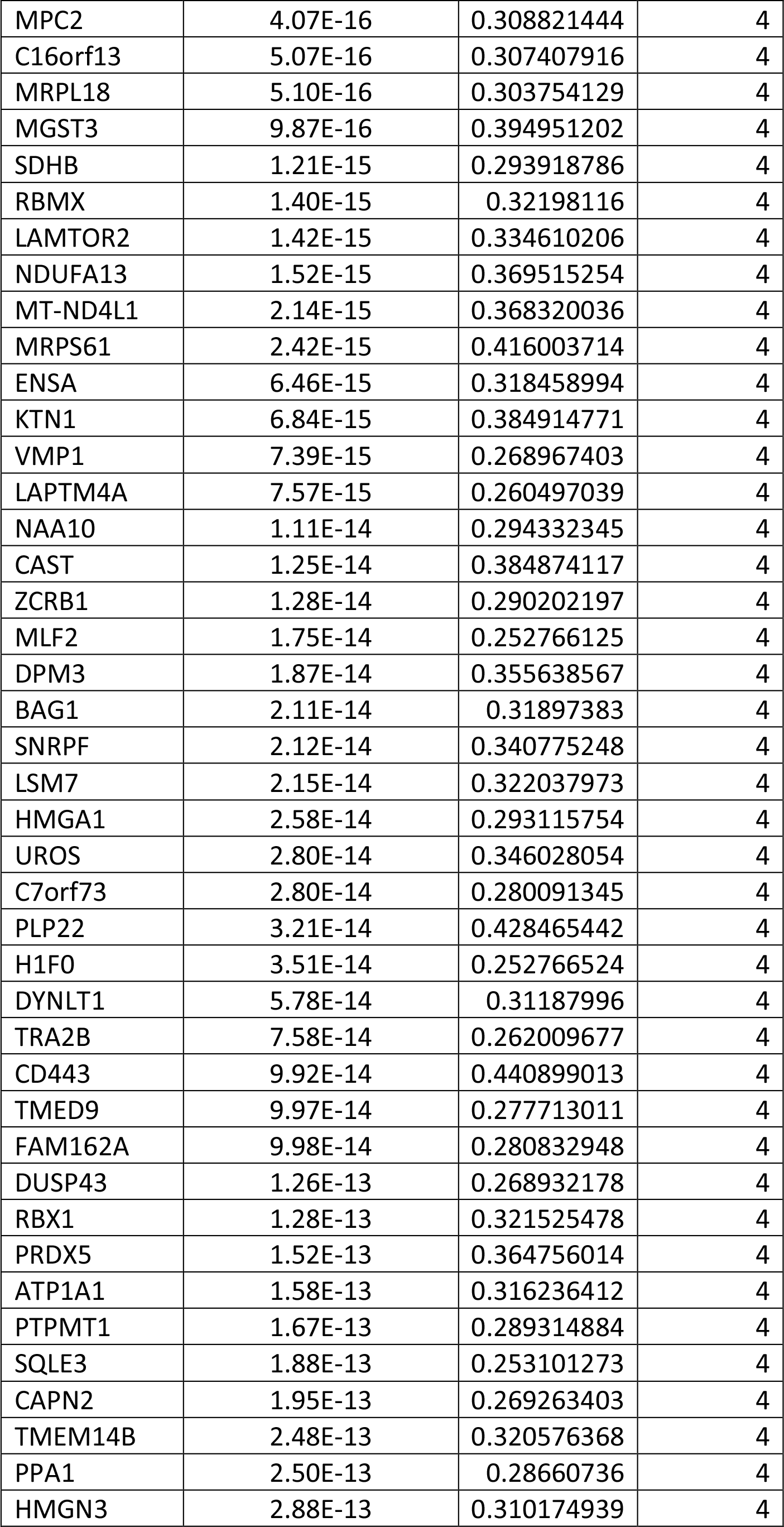
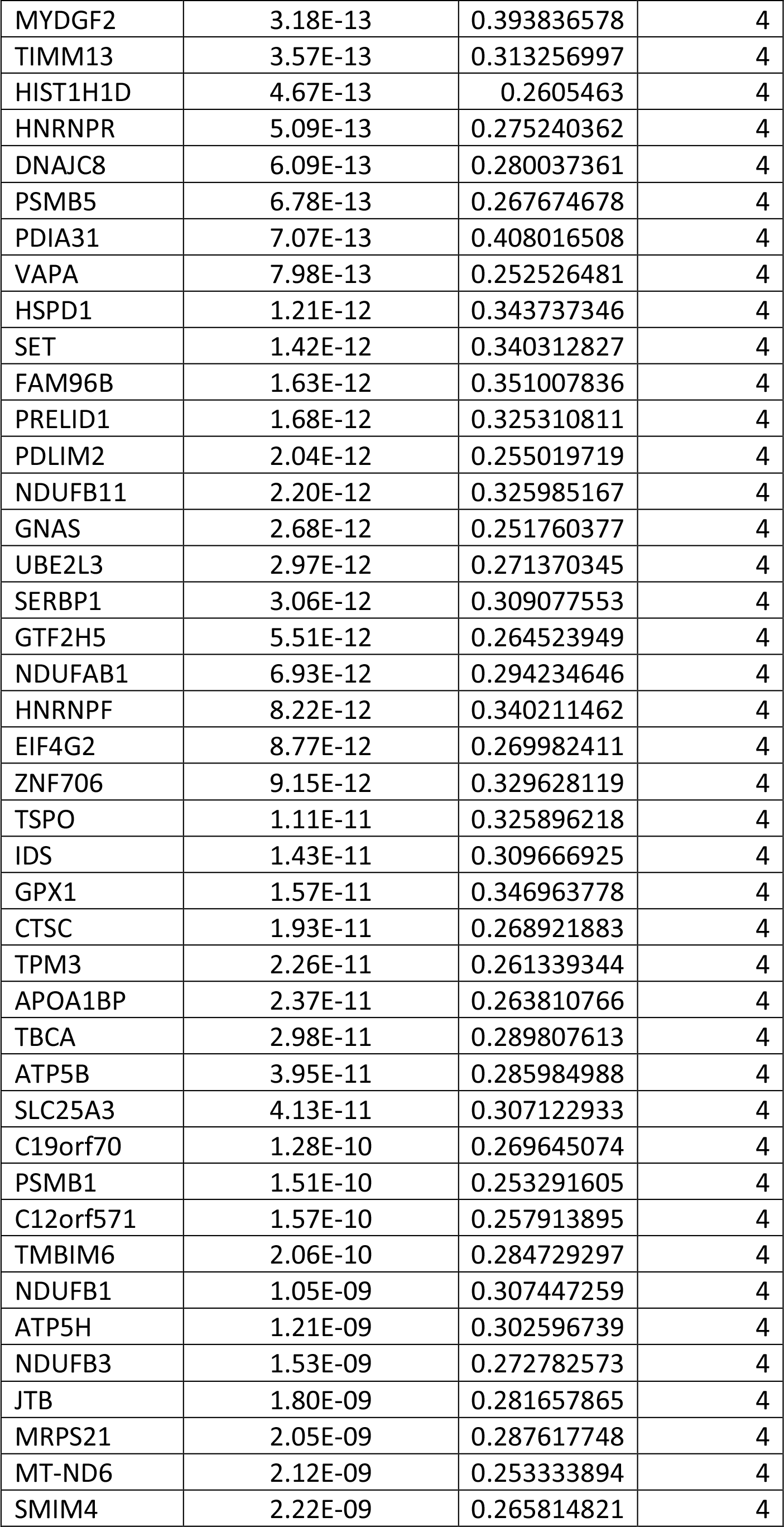
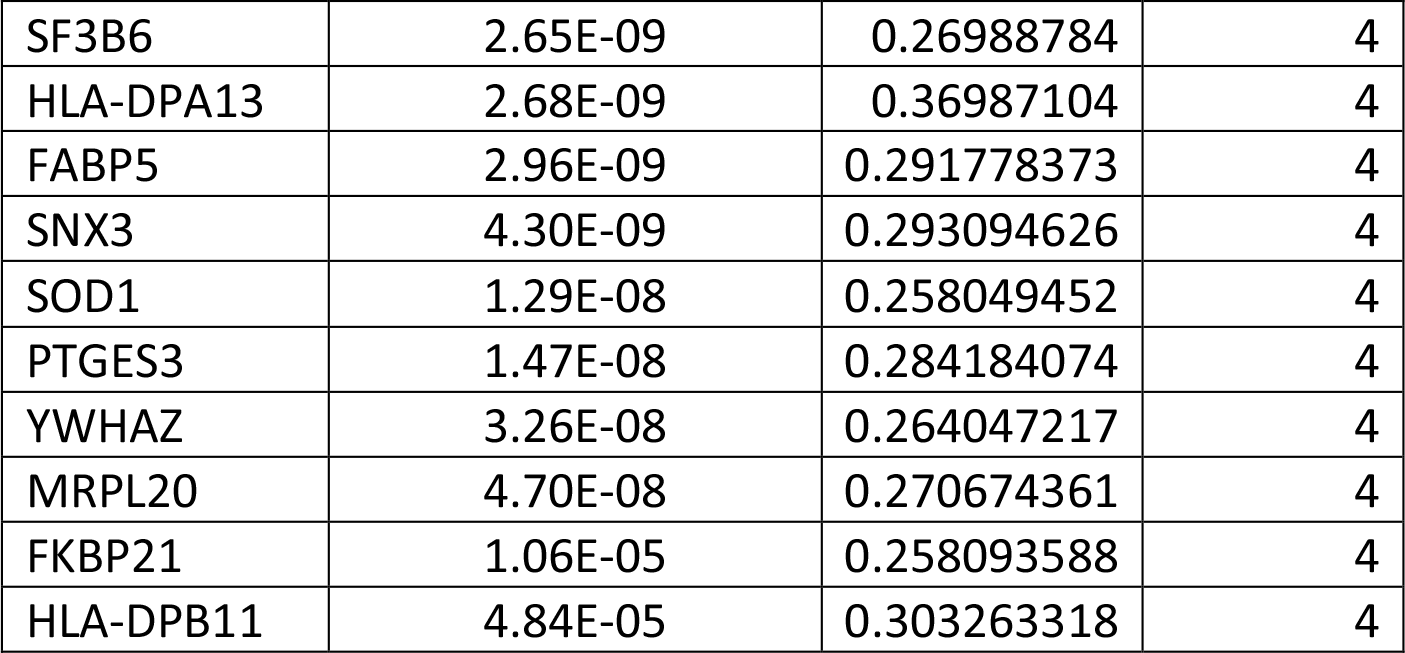
The set of genes that are differentially expressed between clusters of the infected cell population was determined. Genes are ranked by average log fold change. p values were calculated by a Likelihood ratio test.

